# *Solanum* pan-genomics and pan-genetics reveal paralogs as contingencies in crop engineering

**DOI:** 10.1101/2024.09.10.612244

**Authors:** Matthias Benoit, Katharine M. Jenike, James W. Satterlee, Srividya Ramakrishnan, Iacopo Gentile, Anat Hendelman, Michael J. Passalacqua, Hamsini Suresh, Hagai Shohat, Gina M. Robitaille, Blaine Fitzgerald, Michael Alonge, Xingang Wang, Ryan Santos, Jia He, Shujun Ou, Hezi Golan, Yumi Green, Kerry Swartwood, Gina P. Sierra, Andres Orejuela, Federico Roda, Sara Goodwin, W. Richard McCombie, Elizabeth B. Kizito, Edeline Gagnon, Sandra Knapp, Tiina E. Särkinen, Amy Frary, Jesse Gillis, Joyce Van Eck, Michael C. Schatz, Zachary B. Lippman

## Abstract

Pan-genomics and genome editing technologies are revolutionizing the breeding of globally cultivated crops. A transformative opportunity lies in the reciprocal exchange of genotype-to-phenotype knowledge of agricultural traits between these major crops and hundreds of locally cultivated indigenous crops, thereby enhancing the diversity and resilience of our food system. However, species-specific genetic variants and their interactions with desired natural or engineered mutations pose barriers to achieving predictable phenotypic effects, even between closely related crops or genotypes. Here, by establishing a pan-genome of the crop-rich genus *Solanum* and integrating functional genomics and genetics, we show that gene duplication and subsequent paralog diversification are a major obstacle to genotype-phenotype predictability. Despite broad conservation of gene macrosynteny among chromosome-scale references for 22 species, including 13 indigenous crops, hundreds of global and lineage-specific gene duplications exhibited dynamic evolutionary trajectories in paralog sequence, expression, and function, including among members of key domestication gene families. Extending our pan-genome with 10 cultivars of African eggplant and leveraging quantitative genetics and genome editing, we uncovered an intricate history of paralog emergence and evolution within this indigenous crop. The loss of an ancient redundant paralog of the classical regulator of stem cell proliferation and fruit organ number, *CLAVATA3* (*CLV3*), was compensated by a lineage-specific tandem duplication. Subsequent pseudogenization of the derived copy followed by a cultivar-specific structural variant resulted in a single fused functional copy of *CLV3* that modifies locule number alongside a newly identified gene controlling the same trait. Our findings demonstrate that paralog diversifications over short evolutionary periods are critical yet underexplored contingencies in trait evolvability and independent crop domestication histories. Unraveling these contingencies is crucial for translating genotype-to-phenotype relationships across related species.

## INTRODUCTION

Global food production is currently based on fewer than 10 intensively bred commodity crops from only three plant families^1^: grasses (corn, rice, sugarcane, wheat), legumes (soybean), and nightshades (potato, tomato). In contrast, indigenous crops comprise a large, heterogeneous group of hundreds of species which could contribute to agricultural biodiversity and resilience^2^. Many indigenous crops belong to the same families as the major crops but are differentiated by their narrower range of cultivation and scale of production^3^. For instance, the grasses millet (*Eleusine coracana*) and teff (*Eragrostis tef*) and the legumes cowpea (*Vigna unguiculata*) and pigeonpea (*Cajanus cajan*) are locally adapted and important to diets in specific regions of Africa and Asia^4–6^. Within the nightshade (*Solanaceae*) family, the genus *Solanum* alone contains dozens of crop and wild species cultivated in specific regions of Africa and South America for their leaves and/or, fruits, including African eggplant (*S. aethiopicum*), naranjilla (*S. quitoense*), African black nightshade (*S. scabrum*) and pepino (*S. muricatum*)^7,8^.

Indigenous crops are viewed through several different lenses—agricultural, ethnobotanical, and scientific—each with its own unique biases and objectives^2,3,9,10^. Bridging and harmonizing these viewpoints offers an opportunity to better serve local communities and encourage broader adoption for industrialization. Breeding of indigenous crops has been limited relative to global commodity crops. It is widely assumed that decades of research on major crops, along with advances in genome sequencing and genome editing technologies, can be leveraged to address residual undesirable ancestral traits that limit productivity of indigenous, locally adapted crops^11,12^. Engineering beneficial mutations could help rapidly expand the diversity of food species beyond our current genetically narrow, industrialized agricultural systems^2,13^. Despite great progress in genome engineering technologies, however, background dependencies—species-specific genetic modifiers that lead to unpredictable phenotypic outcomes even between closely related species or varieties—remain underappreciated barriers^14^. Indeed, plant breeders have long lamented that beneficial alleles and quantitative trait loci (QTLs) often underperform when transferred to different backgrounds due to interactions among variants^15,16^—a challenge that will persist with genome editing^17,18^.

Our recent tomato pan-genome and associated functional genetics have demonstrated that gene duplications can be potent sources of background modifiers^19,20^. Duplications initially result in genetic redundancy which permits the accumulation of mutations in coding and *cis*-regulatory sequences through genetic drift. Consequently, paralog redundancy can degrade, leading to three canonical outcomes over long evolutionary time: gene loss (pseudogenization), partitioning of ancestral functions (subfunctionalization) or gain of new functions (neofunctionalization)^21,22^. However, the dynamics of how paralogs diverge over shorter time frames, in their sequences, expression patterns, and functions, is less well understood. Genomic and functional dissections of paralogs have largely been limited to within individual species or between widely diverged lineages, and thus have not captured more intermediate trajectories and variable functional consequences of paralog divergence. A deeper understanding of paralog histories and their potentially interdependent relationships could provide greater predictability of phenotypic outcomes when translating genetic knowledge between closely related species. Here, we present a *Solanum* pan-genome and leverage this resource in conjunction with pan-genetics, forward and reverse genetics across species, to comprehensively analyze paralog evolutionary dynamics, demonstrating the value of resolving these underexplored contingencies as we strive to improve indigenous crops for local and climate change adapted agriculture.

## RESULTS

### A chromosome-scale pan-genome of the genus *Solanum*

*Solanum* is one of the most species-rich, ecologically diverse and economically important plant genera^7,8^. The genus includes the major crops eggplant (*S. melongena*), potato (*S. tuberosum*), and tomato (*S. lycopersicum*) and at least 24 indigenous crops, including African eggplant (*S. aethiopicum*), naranjilla (*S. quitoense*) and pepino (*S. muricatum*)^23^. Spanning approximately 16-44 Ma of evolution^24,25^, the diversity of the genus *Solanum*, along with existing genomic and genetic tools in specific species^26,27^, makes it a leading system to study paralog evolution over short evolutionary time scales. We selected 22 species encompassing a broad phylogenetic sample of the ecological (**Fig. 1a**), phenotypic (**Fig. 1b, Extended Data Fig. 1a**), and taxonomic (**Fig. 1c, Supplementary Table 1**) diversity within *Solanum*, including regionally important indigenous crop and ornamental species and several of their wild progenitors. These species are grouped into four main categories that reflect the spectrum of plant use and domestication: wild (W); locally-important, consumed (C); ornamental (O); domesticated food crop (D) (**Fig. 1a,b**). Using PacBio HiFi sequencing and other long-range scaffolding data, we assembled chromosome-scale genomes for all 22 species, including phased haplotypes of the clonally-propagated and highly heterozygous pepino, for a total of 23 assemblies all reaching reference quality (average QV>53, average N50=65.8Mbp) (**Extended Data Fig. 1b,c, Supplementary Table 2**). Final genome sizes ranged from ∼713 Mbp (*S. etuberosum*) to ∼2.5 Gbp (*S. robustum*), with members of the *Lasiocarpa* subclade having four of the five largest genomes. An integrated gene prediction strategy for annotation based on RNA-seq and liftover, allowed us to identify 825,493 high-confidence gene models across the pan-genome (**Extended Data Fig. 1d**, **Supplementary Table 3,** and **Methods**). Of these, 495,429 (∼60%) were shared across all samples, reflecting these species’ relatively close evolutionary relationships.

**Figure 1:**
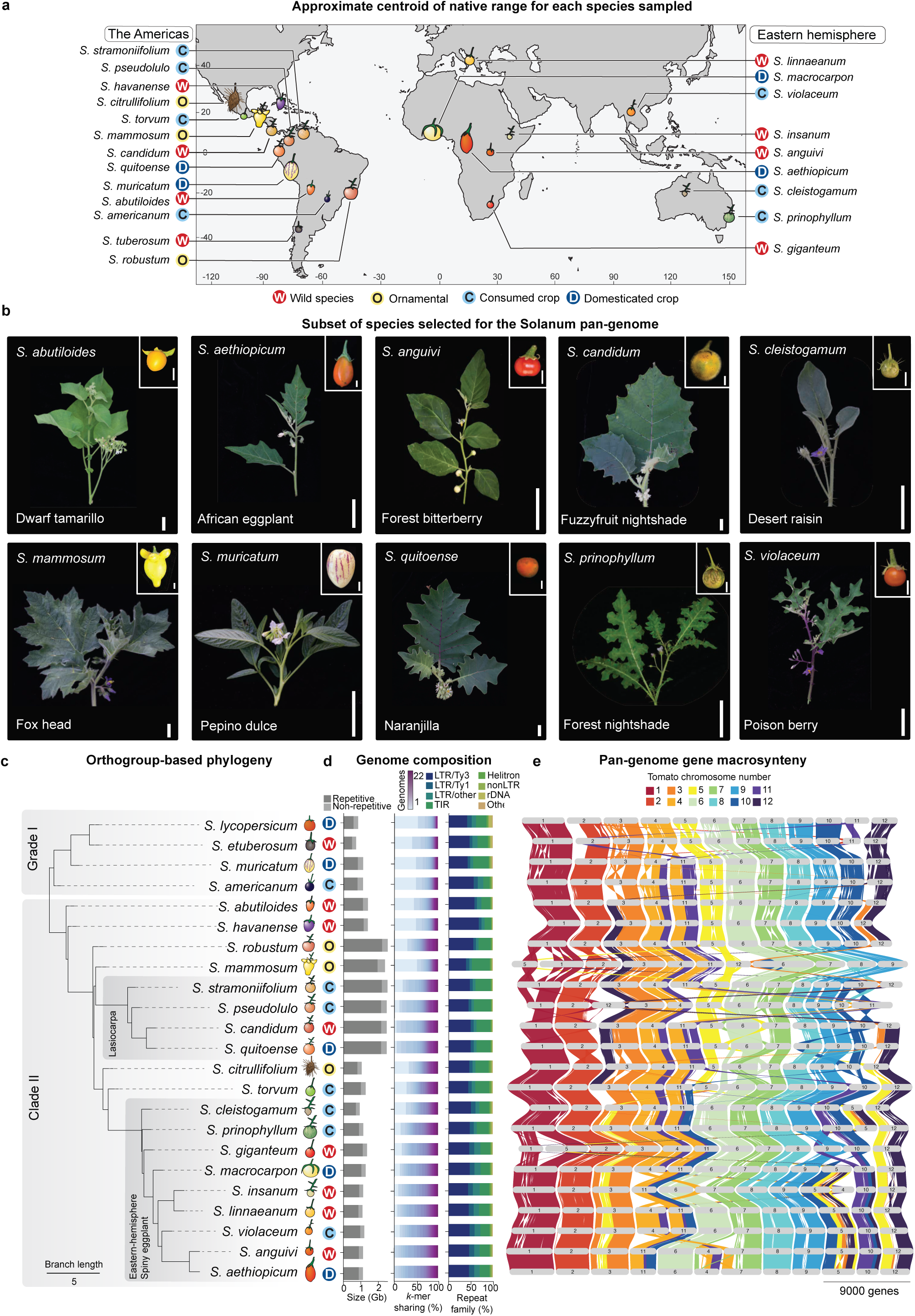
*Solanum* pan-genome captures the phenotypic, ecologic, agricultural, and genomic diversity of this crop-rich genus. **(a)** Approximate centroid of the native range for the 22 selected *Solanum* species, grouped by type of agricultural use: wild (W), locally-important consumed (C), ornamental (O), and domesticated (D). **(b)** Phenotypic diversity of shoots and fruits from a subset of *Solanum* species in the pan-genome. Scale bars: 5 cm (shoots) and 1 cm (fruits). **(c)** Orthogroup-based phylogeny of the *Solanum* pan-genome recapitulates the major clades, Grade I and Clade II. Branch lengths reflect coalescent units. **(d)** Genomic features of each species of the *Solanum* pan-genome. Genome size (Gbp) and representation of non-repetitive (light grey) and repetitive (dark grey) sequences (left). Percentage of pan-k-mers shared across the pan-genome in each reference (middle). Contribution of the different transposable element families in the total repeat landscape of each genome (right). **(e)** GENESPACE plot showing gene macrosynteny across the pan-genome relative to tomato. Scale bar: 9000 genes.

An ortholog-based phylogenetic tree divided the 22 species into two major clades, consistent with previous studies^23,24^. Using existing nomenclature^23^, Grade I included the major crops tomato and potato while Clade II contained all prickly species, including the three cultivated eggplant species: *S. melongena* (Brinjal eggplant), *S. aethiopicum* (African eggplant), and *S. macrocarpon* (Gboma eggplant) (**Fig. 1c**). Whereas gene content was largely uniform across species, transposable element content and distribution varied widely (**Supplementary Table 4**). Consistent with other plant pan-genomes^28,29^, species-specific increases in repetitive content, driven primarily by a rapid expansion of retrotransposon families, correlated strongly with genome size expansion (**Fig. 1d**). The pan-genomic k-mer content – illustrating the genomic diversity within a species relative to the rest of the pan-genome – varied by clade, with 11 species containing more than 25% species-specific sequences (**Fig. 1d**). Finally, ortholog-based analysis revealed broad conservation of gene macrosynteny throughout the pan-genome, with the highest conservation on chromosomes 1, 2, 6, and 9 (**Fig. 1e**). This analysis also revealed large structural rearrangements across the genus and predominantly within sub-clades of clade II, including, for example, megabase-scale inversions and translocations involving chromosomes 3, 5, 10, and 12 (**Fig. 1e**). These high-quality genomes provided a foundation for capturing genetic diversity across the *Solanum* from the clade to the species level, setting the stage for an analysis of paralog evolutionary dynamics and their impacts on genotype-to-phenotype relationships across this species-rich, ecologically and economically important plant genus.

### Pan-genome analysis reveals a complex landscape of gene duplications in *Solanum*

To develop a comprehensive view of gene evolutionary dynamics across *Solanum*, we reconstructed the genus-wide history of orthogroup expansion and contraction events across the 22 species, anchored on tomato (*S. lycopersicum*) **(Fig. 2a)**. From the 44,962 total orthogroups identified across the *Solanum* pan-genome, we identified several of them were involved in expansion (26,284) or contraction (37,267) events, with the majority of the evolutionary events occuring at inner nodes involving orthogroup contractions. Functional enrichment analysis revealed that expanding and contracting orthogroups are predominantly linked to environmental response and secondary metabolism, with species-and clade-specific features (**Extended Data Fig. 2a, Supplementary Table 5, Supplementary Table 6**). We then characterized orthogroups based on their representation in the pan-genome, and classified these orthogroups as core (present in 100% of the species), near core (present in >70% of genomes), dispensable (present in 5-70% of species), and private (found in one species only) (**Fig. 2b**). Most orthogroups are core (60.6%) or near core (20.2%), while smaller proportions are dispensable (14.3%) or private (0.8%). Finally, 75% of pairs of orthologous genes (designated paragroups) are dispensable or private, suggesting derived paralogs are more genetically flexible than orthologs (**Extended Data Fig. 2b**).

**Figure 2:**
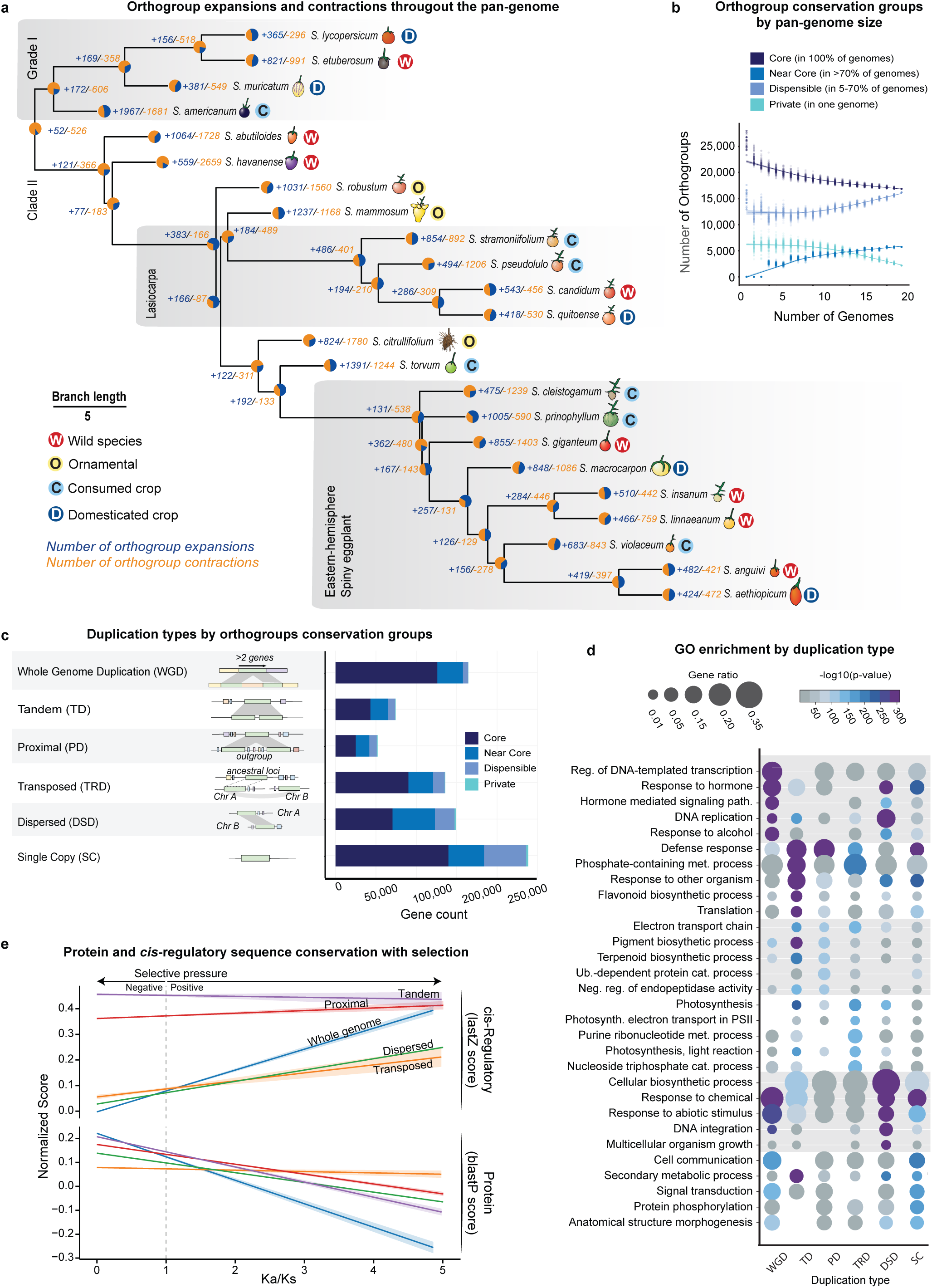
Pan-genomic analysis of orthogroup conservation and diversity of gene duplications. **(a)** Orthogroups expansions and contractions across the pan-genome. The orthogroup-based phylogeny is adapted from Fig. 1c. The estimated expansion (blue) and contraction (orange) rates of orthogroups are shown at each node. **(b)** Cumulative curves showing detection of the four orthogroup conservation groups as a function of the number of species available in the pan-genome. **(c)** Schematic of the potential mechanisms underlying different gene duplication categories (left). Stacked bar chart showing the number of genes derived from the different types of duplication sorted by orthogroup conservation groups (right). WGD: whole-genome duplication; TD: tandem duplication; PD: proximal duplication; TRD: transposed duplication; DSD: dispersed duplication; SC: single copy. **(d)** Functional enrichment of gene duplication types detected across the pan-genome. The top five enriched GO terms per duplication type are shown. Circle size represents gene ratio. **(e)** Divergence of protein and *cis*-regulatory sequences across increasing evolutionary pressure, as measured by Ka/Ks values, for the indicated types of gene duplications. BLASTP (protein sequence conservation) and LASTZ (*cis*-regulatory sequence conservation from the Conservatory algorithm) normalized alignment scores were used to plot the predicted mean and 95% confidence interval.

Across all orthogroups, gene duplications were widespread, with 70% (575,464 duplicates) of all genes having a paralog (**Fig. 2c**). We classified the duplications based on their genomic context as whole-genome (WGD) or single gene duplication, including tandem, proximal, transposed, or dispersed duplications^30^ (**Fig. 2c**). Paralogs most frequently originate from WGDs from events many millions of years ago; however, single gene duplications, which typically are more recent and lineage-specific events, collectively dominate the duplication landscape in *Solanum* (**Extended Data Fig. 2c)**. While most of the WGD-derived duplications belong to core orthogroups, single gene duplications show increased bias towards near core and dispensable orthogroups (**Fig. 2c**). Analysis of duplication types differentiated according to biological function using a GO enrichment analysis show that WGD-derived paralog pairs are most strongly associated with dosage-sensitive processes, such as DNA transcription and DNA replication, as well as hormone-mediated signal transduction and response (**Fig. 2d**), consistent with previous reports^31,32^. In contrast, and as already shown in many systems^30,33^, tandem and proximal duplications are most associated with defense and specialized metabolite biosynthesis, along with diverse functional roles related to environmental responses (**Fig. 2d**).

Paralogous genes functionally diverge through changes in both coding and *cis*-regulatory sequences^34,35^; however, it is unclear if the relative contributions of these changes are associated with specific duplication types. To test this, we first used our previously developed algorithm, Conservatory, which simultaneously allows quantification of *cis*-regulatory conservation and improved calling of paralog pairs based on both protein and *cis*-regulatory conservation^36^ **(Extended Data Fig. 2d** and **Methods)**. We then incorporated Ka/Ks ratios, as a measure of coding sequence selection, with both protein and *cis*-regulatory conservation to determine relationships in coding and regulatory sequence evolution across the duplication types. As expected, for all five types of duplications, protein similarity decreases with higher Ka/Ks values (**Fig. 2e**, **Extended Data Fig. 2e**). However, two striking patterns of *cis*-regulatory conservation distinguish different duplication types: tandem and proximal duplicates maintain high *cis*-regulatory conservation across all levels of selection, whereas WGD, dispersed, and transposed duplicates show higher levels of *cis*-regulatory sequence similarity with increasing Ka/Ks. This observation suggests a greater degree of expression pattern conservation among non-locally duplicated paralogs undergoing functional diversification at the protein level.

### Multi-tissue transcriptomics uncovers the fate of retained paralogs

Research in yeast and other systems suggests that duplicated genes can have negative fitness effects due to increased expression dosage, leading to stoichiometric imbalances in macromolecular complexes^37,38^. Consequently, early diversification of *cis*-regulatory sequences of paralogs may serve to restore ancestral single-copy gene dosage levels in a process called compensatory drift^21,39^. To explore constraints on total expression dosage from retained paralogs, we established two broad categories of paralog pairs as Dosage constrained, or Dosage unconstrained across species and on a per tissue basis (**Fig. 3a**). We defined dosage constrained orthogroups as paralog pairs that exhibited similar total expression levels in a given tissue across species, whereas unconstrained orthogroups did not maintain the same summed expression (**Extended Data Fig. 3a**). To assign paralog pairs to these categories, we generated a pan-*Solanum* gene expression resource comprising 271 samples from 22 species, 15 of which had data from two or more distinct tissues **(Extended Data Fig. 3b**). Principal component analysis (PCA) on the TPM-normalized expression of 5,146 singleton genes showed that the vast majority of samples clustered by tissue type (**Fig. 3b**). As in yeast^40^, our data show that paralog pairs typically evolved under total dosage constraint across tissues and species (**Fig. 3c**). These pairs also exhibited much lower rates of non-synonymous mutations and were less likely to be tissue-specific than unconstrained pairs.

**Figure 3:**
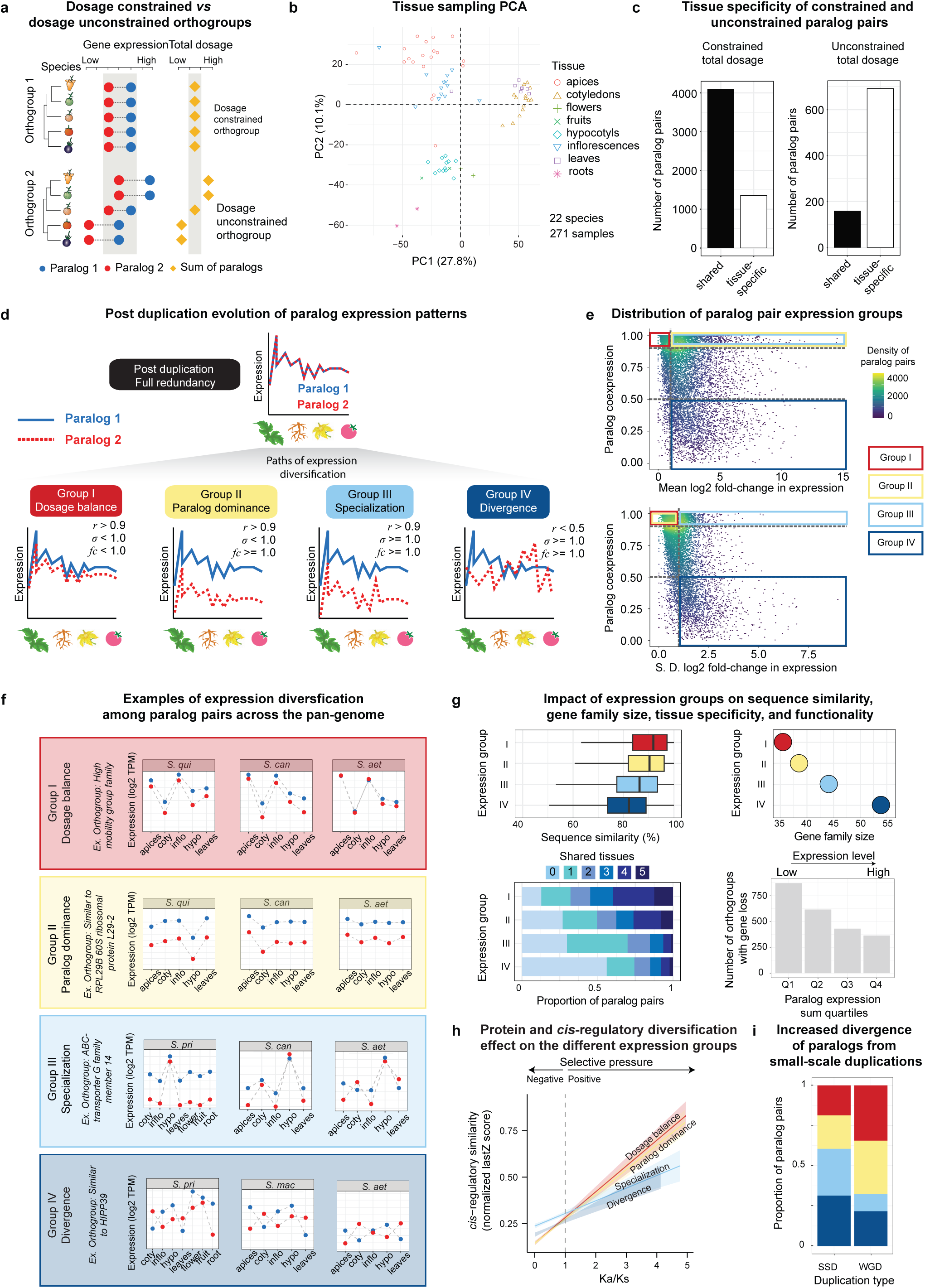
Widespread paralogous diversification across *Solanum* revealed by multi-tissue gene expression analysis. **(a)** Schematic of dosage-constrained and dosage-unconstrained orthogroups reflecting different degrees of selection on the total dosage of paralog pairs across species. **(b)** PCA of the normalized expression matrix from 5,146 singleton genes shared across all 22 species. The expression matrix consists of the summed expression of paralog pairs. Tissue samples are colored by tissue identity. **(c)** Bar plots showing that paralog pairs under constrained total dosage across species are less tissue-specific (left) than unconstrained paralogs (right). **(d)** Schematic of four categories of functional expression groups of retained paralogs: Group I: Dosage balance; Group 2: Paralog dominance; Group III: Specialization; Group IV: Divergence. **(e)** Scatter plots showing the distribution of paralog pairs according to their co-expression level and mean log_2_ fold-change (top) or standard deviation (S.D.) log_2_ fold-change (bottom) in expression. The four derived paralog expression groups are shown. **(f)** Representatives of paralog pairs capturing the different patterns of expression delimited across the pan-genome. **(g)** Genes included in the four paralog expression groups display contrasting protein sequence similarity (top left), gene family size (top right), number of shared expression domains (tissues) (bottom left), or propensity to undergo gene loss for orthogroups in different dosage quartiles (bottom right). **(h)** Effect of *cis*-regulatory sequence conservation on the different expression groups in relation to increased selection on protein sequence. For each expression group the predicted mean and 95% confidence interval of the normalized LastZ score is shown. **(i)** Stacked bar plots showing the proportion of each paralog expression group attributed to paralog pairs derived from either whole-genome duplication (WGD) or small-scale duplication (SSD).

Dosage relationships between paralog pairs can be influenced by different evolutionary trajectories resulting in divergent expression patterns. Among retained paralog pairs within a given species we considered four groups of common patterns of expression relationships following gene duplication (**Fig. 3d**, **Extended Data Fig. 3c**): Group I, Dosage balanced: selection on total dosage remains high, and pairs retain similar expression profiles and levels across tissues; Group II, Paralog dominance: Substantial divergence in expression levels that are proportional across tissues; Group III, Specialization: Expression profiles no longer showing a purely global shift and instead exhibiting tissue-specific changes; Group IV, Divergence: Paralog pairs are fully diverged in both expression profile and level. Applying the definitions to our paralog gene expression dataset showed 58,130 (∼8%) of the paralog pairs to a specific group, leaving over 92% undetermined as they do not yet exhibit strong trajectories **(Fig. 3e,f**, **Extended Data Fig. 3d)**.

While these groups were defined by the expression profiles across tissues within a species, the data also allowed us to evaluate if the groups were associated with distinct genetic features. We compared protein sequence similarity between the groups, as well as gene family function, size, expression status, the number of tissues where expressed, and transcription levels (**Fig. 3g**, **Extended Data Fig. 3e**). We observed that pairs in Group I showed higher sequence similarity, smaller gene family size, broader expression across tissues, and higher transcription levels than groups undergoing paralog dominance, specialization and divergence (Groups II-IV) (**Fig. 3g**). Functional enrichment analysis showed that Groups I-II are enriched in dosage-sensitive processes such as transcription and translation, while Groups III-IV are enriched in defense response genes (**Extended Data Fig. 3e**). Moreover, consistent with their conserved expression patterns, paralog pairs in Groups I and Group II maintained greater *cis*-regulatory sequence conservation than those in Groups III and IV (**Fig. 3h**, **Extended Data Fig. 3f**). We further reasoned that the type of duplications from which gene pairs originated might impact their expression relationships. We found that the most conserved expression groups–paralog pairs in Groups I and II that also capture more ancient duplications–were more likely to have originated from WGDs, whereas gene pairs in Groups III and IV were enriched in small-scale duplications (SSDs) (**Fig. 3i**). Although all four of our defined Groups have the potential to complicate crop engineering, the 60% of pairs with correlated expression patterns likely pose the greatest challenge due to interdependent redundant, compensatory or partially sub-functionalized relationships, which could reflect a continuum of lineage-or specific-specific variations in these relationships.

### Genetic dissection of lineage-specific paralog diversification and compensatory relationships

The *Solanum* pan-genome provided an opportunity to study the extent to which paralog diversifications have shaped key genes that influence genotype-phenotype relationships across the genus. Based on prior characterization and cloning of QTL and developmental genes affecting 16 domestication and breeding traits, we compiled a set of 148 genes and associated paralogs (where relevant) from primarily the three model *Solanum*crops (eggplant, potato, tomato) (**Supplementary Table 7**). Our pan-genome revealed widespread variation in these genes between and within clades, with many cases of gene presence-absence variation (PAV), copy number variation (CNV), and gene truncation/pseudogenization across the pan-genome. Prominent among these were 17 orthogroups containing genes, harboring variants that contribute to the three major components of the crop domestication syndromes (flowering time & plant architecture; inflorescence architecture & flower number; and fruit size) (**Fig. 4a**). For example, in tomato and many other species, variation in the dosage-sensitive florigen-antiflorigen family members (*SP*, *SP5G*, *FTL1a*, *FTL1b*, *SP6D*, *SP6A*, *SFT*) enabled selection for accelerated flowering and short stature (determinate) plants, key traits that facilitated mechanical harvesting^41–43^. We identified numerous CNVs and loss-of-function mutations affecting paralogous genes in our pan-genome, suggesting these variants modulate flowering and growth habit across *Solanum*. In the genetics of inflorescence architecture, mutations in the MADS-box transcription factor-encoding gene *J2* allowed mechanical harvesting of tomato by eliminating the abscission zone of fruit stems^44,45^. However, co-occurring mutations in its ancestral paralog *EJ2* result in undesirable inflorescence branching^46^. We found one CNV and at least three ancestral losses of *J2* in our pan-genome, with most losses occurring in the Eastern Hemisphere Spiny eggplant clade (**Fig. 1c**). These species may therefore be sensitized to changes in inflorescence branching from natural or engineered *EJ2* mutations.

**Figure 4:**
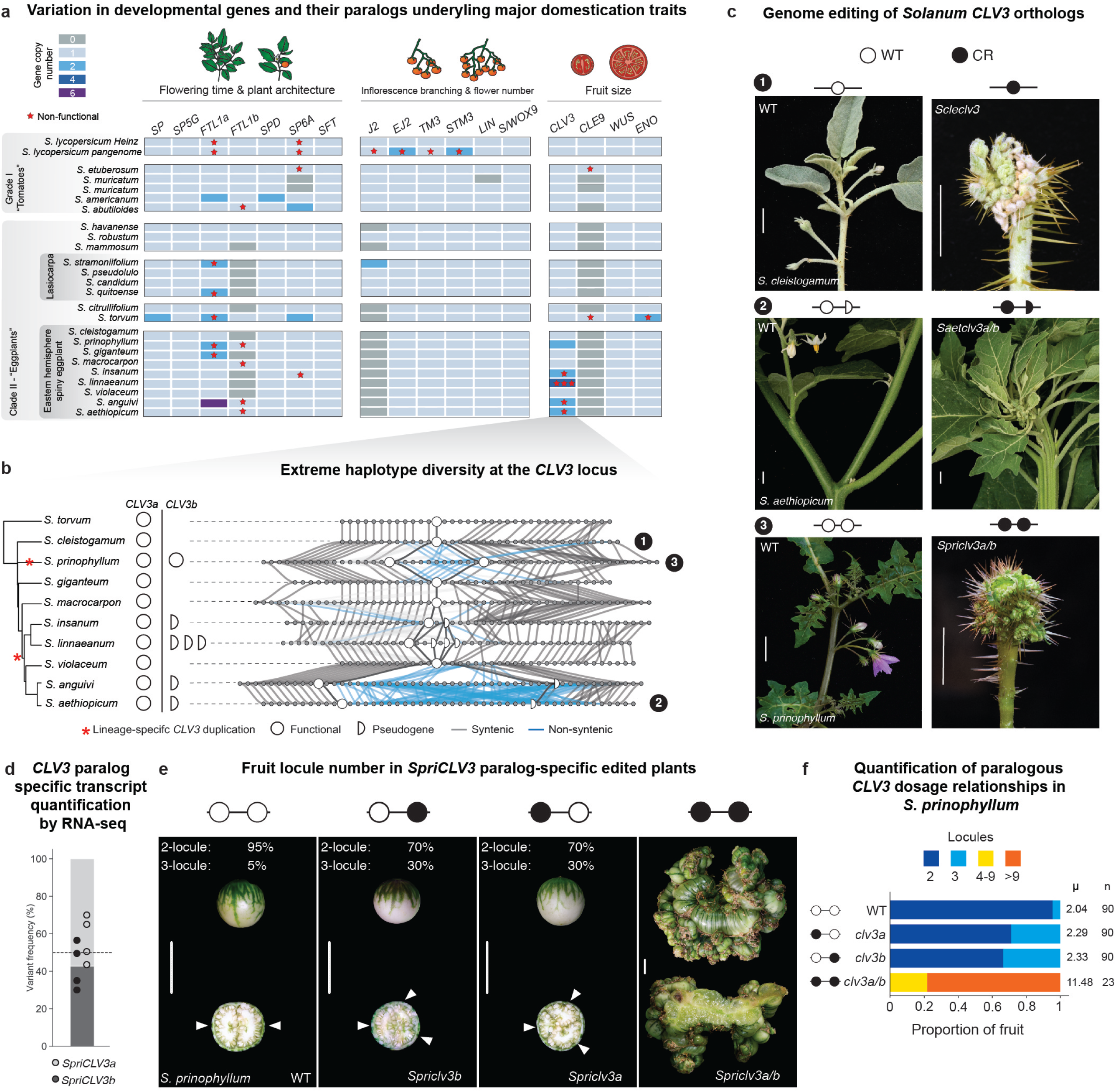
Functional dissection of lineage-specific paralog diversification through pan-genetics reveals modified compensatory relationships in a major fruit size regulator. **(a)** Pan-genome-wide gene presence-absence and copy number variation in 17 orthogroups containing genes known to regulate three major domestication and improvement traits in tomato. Stars indicate gene truncation or pseudogenization. **(b)** Haplotype diversification at the *CLV3* locus across the eggplant clade. Presence-absence of *CLV3* paralogs is shown. Lineage-specific *CLV3* duplications are marked with asterisks. Full circles denote functional *CLV3* copies and half circles denote truncated/pseudogenized copies. Grey lines illustrate conservation, while blue lines represent loss of synteny. **(c)** CRISPR/Cas9 genome editing of *CLV3* orthologs in three species of the eggplant clade. Engineered loss-of-function mutations in *S. cleistogamum* (*ScleCLV3*, top), *S. aethiopicum* (*SaetCLV3a/b,* middle), and *S. prinophyllum* (*SpriCLV3a/b,* bottom) resulted in severely fasciated stems and flowers in all three species. Scale bars: 1 cm. **(d)** Quantification of *SpriCLV3* paralog-specific transcripts by RNA-seq. **(e)** Locules per fruit after paralog-specific gene editing of *SpriCLV3a* and *SpriCLV3b* in *S. prinophyllum*. Single paralog mutants cause a subtle shift from bilocular to trilocular fruits; inactivation of both paralogs results in highly fasciated fruits. Arrowheads mark locules. Scale bars: 1 cm. **(f)** Quantification of locule number in single and double *Spriclv3a* and *Spriclv3b* mutants. Proportion of each locule number per genotype is shown.

The increase of fruit size in tomato domestication was driven in large part by a promoter structural variant in the stem-cell signaling peptide gene, *CLAVATA3* (*CLV3*)^47^*. CLE9,* a partially redundant ancestral paralog, falls into Group II (paralog dominance) and partially compensates for the effect of the *CLV3* domestication allele^48,49^. We previously showed *CLE9* was pseudogenized or completely lost in several Solanaceae species, which eliminated partial redundancy with *CLV3*^48^. Notably, except for tomato and *S. americanum,* all species in our pan-genome contain a pseudogenized *CLE9* or lack it entirely. Meanwhile, a subset of the Eastern Hemisphere Spiny eggplant clade possess locally duplicated intact and pseudogenized copies of *CLV3* (**Fig. 4a, b**). Our chromosome-scale references revealed complex haplotypes involving these duplications, with species-specific transposable element invasions and disease resistance genes interspersed between the paralogs. For example, whereas *S. prinophyllum* carries two intact copies of *CLV3,* one intact and one to three pseudogenized copies exist in *S. aethiopicum* (African eggplant, 1 pseudogenized copy), its progenitor *S. anguivi* (1 pseudogenized copy), and *S. linnaeanum* (3 pseudogenized copies), with extreme variation in transposable element and disease resistance gene content and structure (**Fig. 4b**, **Extended Data Fig. 4a,b**). Comparing haplotypes and observing identical breakpoints in pseudogene structure across these species suggested at least two independent *CLV3* duplication events in the Eastern Hemisphere Spiny clade where one ancestral duplication was followed by pseudogenization in the last common ancestor of *S. insanum, S. linnaeanum, S. anguivi,* and *S. aethiopicum,* whereas a more recent *CLV3* duplication emerged in the lineage leading to *S. prinophyllum* (**Fig. 4b)**. However, we cannot exclude the possibility of three independent duplications, as *S. violaceum* carries only one *CLV3* copy.

The independent duplication resulting in two intact copies of *CLV3* in *S. prinophyllum* suggests redundancy was re-established in this species (Group I), whereas in species where one *CLV3* paralog was pseudogenized, redundancy was again lost. We tested this by using CRISPR/Cas9 to inactivate *CLV3* in three spiny *Solanum* species: *S. cleistogamum* (desert raisin – *ScleCLV3* single copy), *S. aethiopicum* (African eggplant – one functional (*SaetCLV3a*) and one pseudogenized (*SaetCLV3b*)), and *S. prinophyllum* (intact copies of *SpriCLV3a* and *SpriCLV3b*) (**Fig. 4c**, **Extended Data Fig. 4c,d**). As expected, mutations in the one intact copy of *CLV3* in *S. cleistogamum* and *S. aethiopicum* resulted in extreme fasciation phenotypes, matching tomato *clv3 cle9* double mutants (**Fig. 4c**). Similarly, knocking out both copies of *CLV3* in *S. prinophyllum* (*SpriCLV3a* and *SpriCLV3b*) replicated this severe phenotype.

*SpriCLV3a* and *SpriCLV3b* in *S. prinophyllum* are identical in their coding and *cis*-regulatory sequences, except for a single nucleotide variant in the 3’ untranslated region (UTR) of the ancestral copy. Such high sequence identity suggested that eliminating one copy would be fully compensated for by the remaining functional copy, similar to the near complete compensation between *PgriCLV3* and *PgriCLE9* in the *Solanaceae* species *Physalis grisea* (groundcherry)^48^. Our previously generated transcriptomic data of meristems from *S. prinophyllum*^50^ showed both paralogs are expressed to similar levels (**Fig. 4d)** and supported this prediction. Surprisingly, we found that engineered mutations in either *SpriCLV3* paralog resulted in a subtle shift to more trilocular fruits compared to wild type (5% trilocular in WT compared to 30% trilocular in single mutants), suggesting one paralog cannot fully compensate for the other, most likely because of a gene expression dosage effect (**Fig. 4e,f, Supplementary Table 8**).

Taken together, these data suggest that, following the loss of the ancestral *CLE9* paralog, subsequent tandem duplication events in three spiny *Solanum* lineages would have reestablished *CLV3* compensation. However, this compensation was then lost again in at least one lineage due to pseudogenization of the derived *CLV3* duplicate. Finally, despite retention of both nearly identical copies of *CLV3* in *S. prinophyllum*, complete compensation was not fully maintained. Similar to *CLV3*, dynamic duplication histories and resulting paralog relationships affecting meristem proliferation and other gene families critical for domestication and trait improvement may reveal the species-specific contingencies that impact outcomes in genome engineering.

### African eggplant pan-genomics reveals widespread introgression and paralog diversification

African eggplant (*S. aethiopicum*) is a major crop indigenous to sub-Saharan Africa and cultivated across the continent on hundreds of thousands of acres. Transported by enslaved Africans, it is also grown extensively in Brazil, but outside of these regions it remains largely unknown (**Fig. 5a**)^51,52^. Diverse cultivars are grown in Africa for their edible fruits or leaves, as well as for the ornamental appeal of specific fruit types^53^. These disparate uses are reflected in the species’ broad intraspecific diversity in vegetative and fruit phenotypes, including fruit shape, color, and size (**Fig. 5b**). Breeding in African eggplant has primarily focussed on improving adaptation to abiotic stress conditions^54,55^, with less progress on improving yield or productivity. Re-engineering or mimicking the effects of known beneficial mutations from tomato and other *Solanum* model crops could advance these goals, but genomic and genetic resources are limited.

**Figure 5:**
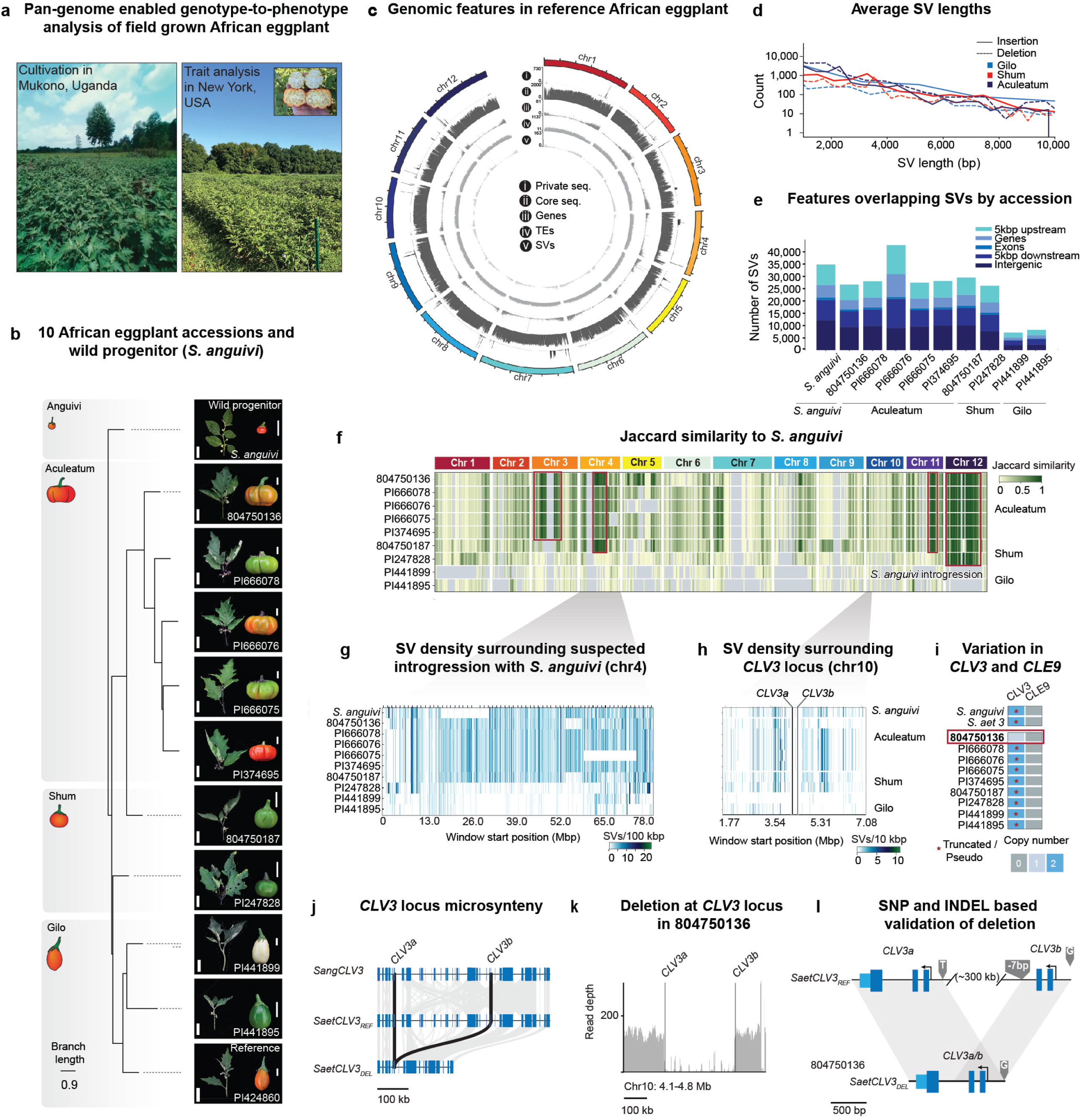
Pan-genome of African eggplant reveals widespread structural variation, wild species introgression, and *CLV3* paralog diversification. (a) Images of field-grown African eggplant in Mukuno, Uganda (left) and New York, USA (right). **(b)** Ortholog-based phylogeny of 10 African eggplant accessions covering three main cultivar groups (Gilo, Shum, and Aculeatum) and the wild progenitor *S. anguivi*. Representative shoots and fruits are shown for each accession. Scale bars: 5 cm (shoots), 2 cm (fruits). Branch lengths reflect coalescent units. **(c)** Pan-genomic features across African eggplant reference genome. Frequencies of: (i) sequences private to the reference, (ii) core sequence, (iii) genes, (iv) transposable elements, and (v) SVs. **(d)** Average SV lengths (bp) for deletions (dotted lines) and insertions (solid lines) across the three African eggplant cultivar groups. **(e)** Number of SVs overlapping with genomic features across accessions. **(f)** Jaccard similarity of SVs across the African eggplant pan-genome measured against *S. anguivi* in 2 Mbp windows. Putative introgression from *S. anguivi* on chromosomes 3, 4, 11, and 12 are highlighted by red boxes. **(g)** Close-up of chromosome 4 introgression shown by SV density. **(h)** SV density surrounding the *SaetCLV3* locus across the pan-genome. Genomic positions of *SaetCLV3a* and *SaetCLV3b* are shown. Window size: 10 kbp. **(i)** Presence-absence and copy number variation of *CLV3* across the pan-genome. *CLE9* is absent in all genotypes. *S. aethiopicum* and *S. anguivi* are shown for reference. **(j)** Conservation of exonic microsynteny (grey bars) between *SangCLV3*, *SaetCLV3_REF_*, and *SaetCLV3_DEL_* haplotypes. Scale: 100 kb. **(k)** Long-reads pile-up at the *SaetCLV3* locus identifies a deletion structural variation and distinct *SaetCLV3* haplotype in accession 804750136. **(l)** Diagram of deletion-fusion allele of *CLV3* (*SaetCLV3_DEL_*) arose in accession 804750136. The 7 bp indel and SNPs were used as markers to validate the deletion-fusion scenario.

To address this, we first phenotyped eight representative accessions (**Supplementary Table 9**) from the Gilo (fruit production), Aculeatum (ornamental), and Shum (leaf production) cultivar groups in field conditions (**Fig. 5a**) along with one accession of *S. anguivi*. Based on the observed phenotypic variation, we extended our selection to 10 diverse accessions belonging to the three cultivar groups (**Supplementary Table 9**) and assembled a long-read based African eggplant pan-genome that included its wild progenitor *S. anguivi*. The reference African eggplant accession (PI 424860) belongs to the Gilo group, and was used as the representative genotype in the wider *Solanum* pan-genome (**Fig. 1**). To assess genetic relationships, we computed an ortholog-based phylogenetic tree (**Fig. 5b**), which indicated two major clades, one comprising the three Gilo accessions and a second containing the five Aculeatum accessions. Interestingly, the two Shum accessions did not form a monophyletic group, suggesting that accessions cultivated for leaf production might have different genetic origins. Protein-coding genes were primarily clustered at chromosome ends throughout the African eggplant pan-genome, a pattern similar to other *Solanum* and flowering plant species (**Fig. 5c**). Transposable element distribution complemented this pattern, with more elements accumulating in the gene-poor pericentromeric regions.

Comparing the African eggplant genomes against the reference showed that, at the sequence level, most of the genome is highly conserved. Over 250,000 structural variants (SVs: defined as variants at least 50 bp in size) were found across all African eggplant samples, mainly towards chromosome ends (**Fig. 5d**, **Extended Data Fig. 5a**). Similar to our tomato pan-genome^19^, over 68% of SVs were located within 5 kbp upstream or downstream of genes, in addition to 7,234 SVs overlapping exons and therefore likely to disrupt gene function (**Fig. 5e**, **Extended Fig. Data 5b**). While average SV length was similar across accessions, their absolute number varied between groups, with Gilo possessing the fewest SVs, an expected pattern since the reference African eggplant belongs to the Gilo group. Notably, the SV distribution showed clade-specific SVs and SV clusters shared with the wild ancestor *S. anguivi,* suggesting a history of introgression (**Fig. 5f**). Using a window-based Jaccard similarity analysis, we found multiple introgressions from *S. anguivi* in the Aculeatum accessions, most evident on chromosomes 3, 4, 11, and 12. Such widespread introgression suggests recent gene flow from the wild species in the course of African eggplant breeding, and likely explaining the origin of the Aculeatum ornamental types (**Fig. 5b, f, g**).

Similar to tomato, African eggplant cultivar groups exhibit extreme variation in fruit size, based in large part on variation in locule number (**Fig. 5b**). We reasoned that, beyond interspecific paralog dynamics observed throughout the pan-genome, recent diversification of key regulators of fruit locule number, such as *SaetCLV3*, might have favored intraspecific phenotypic diversity. The *SaetCLV3* locus, located on chromosome 10, is nested in dense SV clusters (**Fig. 5h**). Interestingly, one Aculeatum accession (804750136) has only a single intact copy of *SaetCLV3,* suggesting the ancestral pseudogenized copy was eliminated (**Fig. 5i**, **Extended Data Fig. 5c**). Microsynteny analysis revealed broad rearrangements at the *CLV3* locus between African eggplant and *S. anguivi*, as well as intraspecific diversity (**Fig. 5j**). Notably, we detected two deletions within the *SaetCLV3* locus in two *S. aethiopicum* accessions (804750136 and PI 247828), including a ∼300 kbp deletion between the second exon of *SaetCLV3a* and the first exon of *SaetCLV3b* (**Fig. 5k**). Remarkably, the large deletion resulted in a fusion between the intact and pseudogenized *SaetCLV3* copies, resulting in a single functional copy, designated *SaetCLV3^DEL^* (**Fig. 5k**).

### Paralog contingencies impact fruit locule number step changes in African eggplant

We next sought to understand if *SaetCLV3* haplotype and paralog dynamics influenced locule number variation. Using our African eggplant genomes, we performed QTL-seq to map loci controlling locule number (**Supplementary Tables 10, 11, 12**). We generated F2 mapping populations between the high-locule count reference accession (PI 424860) belonging to the Gilo group and low-and high-locule count parents belonging to the Shum (804750187) and Aculeatum (804750136) groups, respectively (**Fig. 6a**, **Extended Data Fig. 6a**). In contrast to tomato, the major step change in locule number between the Gilo and Shum groups mapped to a QTL in a 3.9 Mbp region on chromosome 2, which conspicuously did not include *CLV3* or any other known *CLV* pathway components (**Fig. 6b**). Instead, we identified a candidate gene encoding a serine carboxypeptidase (*SaetSCPL25-like*, named after its best BLAST hit in *Arabidopsis*^56^) harboring a 5 bp exonic frameshift deletion in the Gilo parent. Serine carboxypeptidases function in C-terminal peptide processing, and such control of CLE peptide processing has been demonstrated in *Arabidopsis*, where mutation of the Zn^2+^ carboxypeptidase-encoding gene *SOL1* (*Suppressor of LLP1*) represses CLE-dependent root meristem size-related defects^57^. The mutation in *SaetSCPL25-like* in African eggplant was associated with the development of approximately two additional locules (**Fig. 6c**). We validated this association by mutating the orthologs of this gene in both tomato and *S. prinophyllum* using CRISPR/Cas9, which caused quantitatively similar locule number increases as the natural mutation in African eggplant **(Fig. 6d)**.

**Figure 6:**
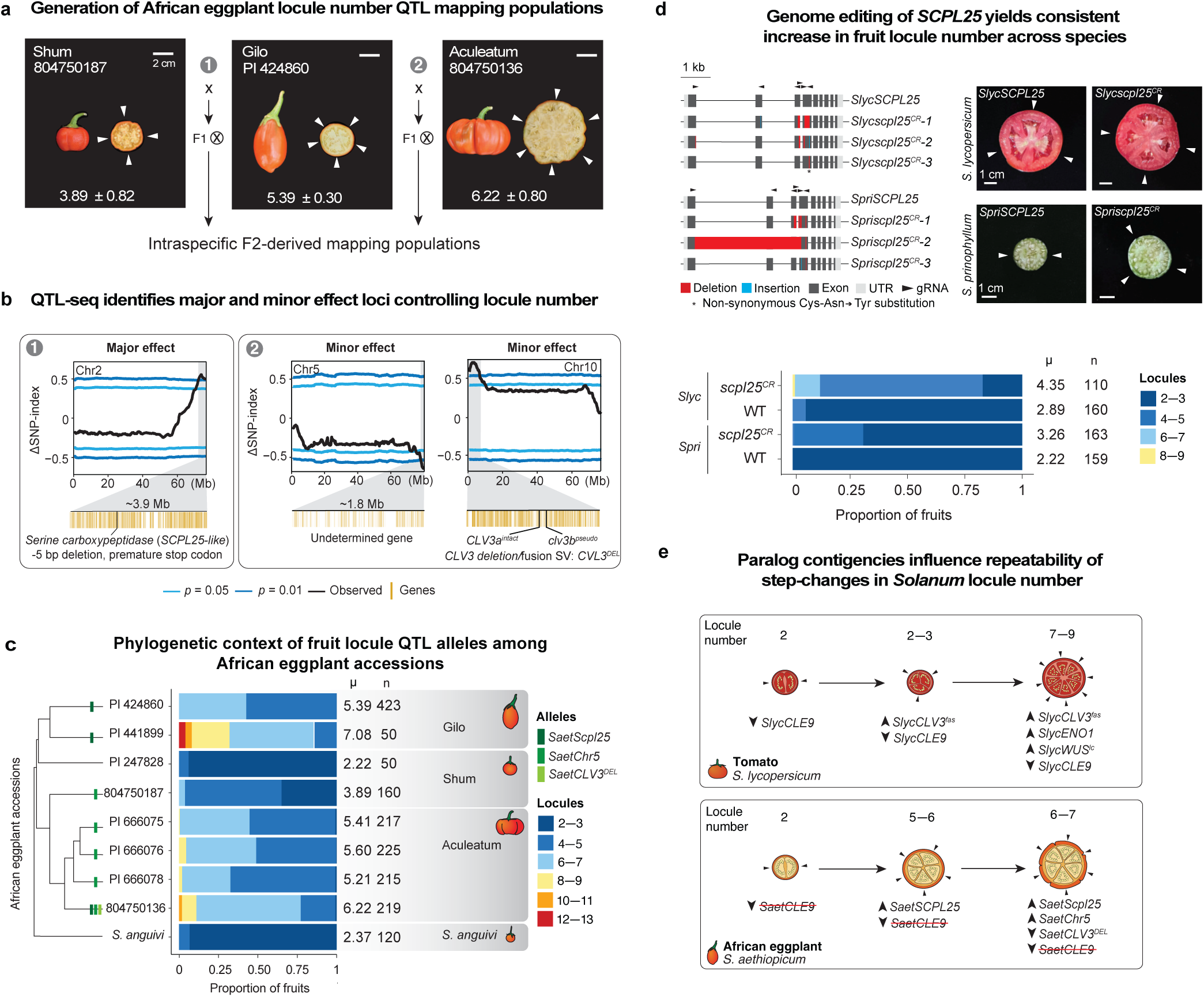
Pan-genetic dissection of fruit locule variation in African eggplant. (**a**) Intraspecific crosses between representative accessions of each of the three main cultivated groups of African eggplant were used to generate F2 mapping populations for QTL-Sequencing (QTL-seq). (**b**) Major (1) and minor (2) effect QTLs affecting locule number identified by bulk-segregant QTL-Seq. ΔSNP-indices for three identified QTL on chromosomes 2, 5, and 10 indicate the relative abundance of parental variants in bulked pools of F2 individuals (low and high locule classes) calculated in 2000 kbp sliding windows. **(c)** Stacked bar plots showing fruit locule number from phylogenetically-arranged African eggplant accessions. Presence of the three mapped QTL alleles (different intensity green bars) in each accession are indicated on the phylogenetic tree. **(d)** CRISPR/Cas9 engineered mutant alleles of *SCPL25* serine carboxypeptidase orthologs in tomato (*SlycSCPL25*) and *S. prinophyllum* (*SpriSCPL25*) (left), along with representative images of transverse fruit sections from mutant plants (right) and quantification of fruit locule number (bottom). Scale bars: 1 cm. **(e)** Schematics comparing the genetic basis of step changes underlying increased locule number and fruit size in tomato and African eggplant. Arrowheads in transverse fruit depictions indicate locules. Average fruit locule number (μ) and fruit number (n) are indicated to the right of stacked bar plots.

We also identified two minor effect QTLs from the Aculeatum group, which we mapped to a 1.8 Mbp region on chromosome 5 and a 4.9 Mbp region on chromosome 10. The latter encompasses the *SaetCLV3^DEL^* haplotype harboring the reconstituted single copy functional *SaetCLV3* (**Fig. 5e** and **Fig. 6c**). We found that these two minor effect QTLs interact, with the homozygous *SaetCLV3^DEL^* genotype masking the increase in locule number conferred by the chromosome 5 haplotype derived from the Aculeatum parent (**Extended Data Fig. 6b**). Though the specific gene and variant underlying the chromosome 5 QTL remains to be characterized, these results indicate that multiple interacting loci, two of which affect the CLV3 signaling pathway, gave rise to increases in locule number in African eggplants.

We then asked how these QTLs shaped the domestication history of African eggplant by examining the alleles present at the three identified loci within the phylogenetic context of our African eggplant pan-genome (**Fig. 6c**). The Gilo accessions contained the *SaetSCPL25-like* mutant allele, while all surveyed Aculeateum accessions and one of the Shum accessions harbored the chromosome 5 minor effect QTL’s haplotype. Meanwhile, a single Aculeatum accession (804750136) contained all three identified alleles, including the minor effect *SaetCLV3^DEL^*structural variant (**Fig. 6c**). The SV at *SaetCLV3* probably occurred secondarily to variants at *SaetSCPL25-like* and the chromosome 5 QTL. *SaetCLV3^DEL^*causes a subtle reduction in locule number, and was perhaps selected to attenuate the locule count increases conferred by the combined synergistic effect of *SaetSCPL25-like* and the chromosome 5 QTL (**Extended Data Fig. 6b)**. This contrasts with tomato, where previous studies identified the *SaetCLV3* structural variant *SlycCLV3^fas^* as a widespread and major effect QTL variant yielding increased fruit locule number, modified by other minor effect QTLs, including the paralog *SlCLE9*^49^. Thus, while QTLs affecting *CLV* signaling are shared drivers of increased locule number in both tomato and African eggplant, the specific genes, alleles, and interactions, as well as the magnitude and directionality of these individual and combined effects, are distinct (**Fig. 6e**). The recurrence of QTLs at *SaetCLV3* in two independent domestication histories underscores the major contribution of structural variation on paralog evolutionary dynamics as key contingencies shaping parallel trajectories of crop domestication and improvement.

## DISCUSSION

Plant pan-genome resources are emerging at an incredible pace. A widespread assumption is that implementing genome editing technologies on these foundational resources will be the panacea to translating genotype-to-phenotype knowledge between related crops and also their wild relatives^11,12^. Decades of work by plant breeders demonstrates, however, that additive and epistatic effects from background genetic modifiers are a barrier to predicting desirable outcomes^14–16,58^. While sequencing high-quality plant references at scale, including potentially telomere-to-telomere genomes^59^, combined with forward genetics, can readily uncover background variation, identifying orthologs and paralogs and tracing their evolutionary trajectories remains an unsolved challenge, particularly given the exceptionally complex history of ancient whole-genome duplications and more recent smaller-scale duplications across flowering plants. This is especially problematic in pan-genomes spanning broader taxonomic scales, where more extreme amounts of sequence variation are found.

We approached the challenge of resolving orthologs, paralogs, and their diversification histories using an integrated approach. We used existing tomato and eggplant annotations, multi-tissue RNA-seq annotations, and manual curation to expose and compare ancient paralogs and recent tandem duplications across our pan-genome. We mapped core and dispensable genes in the pan-genome, and among the tens of thousands of paralog pairs identified, expression analyses revealed a continuum of redundancy relationships, driven by drifting expression patterns, pseudogenization, or gene loss. Most dramatically, we showed that paralogs of the fruit size gene *CLV3* captured all three possible scenarios, reflected in independent tandem duplication events, extreme haplotype shuffling, and pseudogenization, accounting for variation in this domestication trait within and between species. Our approaches showcase how leveraging knowledge from major crops to indigenous crops and wild species can reveal previously unknown factors involved in trait variation, opening the door to reciprocal knowledge gain and new paths to improving all crop species.

Similarly complex paralog evolutionary histories undoubtedly affect other traits in nightshades, grasses, legumes, and beyond. Assembling widely and deeply sampled species and genotypes into super pan-genomes^29,60^ offers watershed opportunities to both better understand origins and frequencies of genome fragility within and between species and mobilize advances in machine learning for *de novo* genetic and genomic predictions at scale. As more accurate machine learning models are developed, the micro-level analysis (e.g. read-level basecalling^61^ or variant detection^62^) as well as higher level predictions of epigenomic and regulatory activity^63^ have been and will be greatly improved and expedited. Efforts to predict gene expression changes from *cis*-regulatory variations are also maturing, although limitations in the modeling frameworks and their training regimes remain obstacles to achieving high predictive accuracy^64^. Advancing these efforts to predict trait variation from both coding and *cis*-regulatory variations will undoubtedly be even more challenging. Our work here shows that such models must explicitly account for paralogs and their diversification dynamics over both short and long evolutionary times. Nevertheless, our ability to predict genotype-to-phenotype relationships, a holy grail for genetics and biology, will inevitably be enhanced by developing a foundation model trained on ever-increasing catalogs of molecular, cellular, and organismal data within and across species, to aid in both plant breeding and understanding natural diversity.

We also recognize that real-world implementation of pan-genomic and pan-genetic resources, tools, and technologies requires a deeper understanding of, and sensitivity to, the central role that indigenous knowledge and cultures have played in botany and agriculture^10,65,66^. Within this project, ethnobotanical knowledge from local breeders provided essential expertise in choosing the lineages, species, and cultivars to give our pan-genome immediate impact in agriculture. This includes the potential to rescue traits of agronomic interest that may have been lost during domestication, such as stress resistance and specialized metabolism^67,68^. This is most exemplified through the inclusion of African eggplant, one of the most economically and culturally important crops in tropical sub-Saharan Africa. Our integrated genomic, transformation, and genome editing pipeline complements the rich genetic and phenotypic diversity available in the African eggplant germplasm, offering new and more predictable avenues for breeding. For example, from dissecting the parallel, but distinct, genetic and epistatic paths towards increased locule number in tomato and African eggplant, we have more power to predictably increase locule number, fruit size and yield in this indigenous and regionally important crop.

We expect additional advancements will come from resolving paralog histories and relationships of flowering regulators, which have been central to agricultural revolutions^69^. It is important to highlight, however, that while industrialized breeding emphasizes yield, the needs of subsistence farmers can be different^70^. In the case of African eggplant, modifying flowering time and inflorescence architecture are arguably as important as increasing fruit size. In varieties grown for fruit production, earlier flowering and more branched genotypes would simultaneously dwarf plants and accelerate fruit production and total yield, whereas in varieties cultivated for leaf consumption, late flowering would prolong vegetative growth and vegetative yield^71,72^. We propose that the florigen-antiflorigen flowering hormone system and its MADS-box gene targets should be the primary targets to achieve these goals. In particular, our study revealed distinct diversifications in African eggplant of both florigen and antiflorigen paralogs from tomato, where there is already deep knowledge of these genes and their functional relationships^69^. Knowledge of these paralogs, their allelic diversity, and epistatic relationships and contingencies will provide opportunities to accelerate breeding of these traits in African eggplant with natural alleles of these genes, which can now be characterized through pan-genome-enabled quantitative genetics, and will facilitate predictable outcomes from genome engineering. Looking forward, the most promising strategies for improving indigenous crops can only be realized through effective communication, understanding, and collaboration among local people, scientists, breeders, and growers.

### Online Content

Any methods, additional references, Nature Portfolio reporting summaries, source data, extended data, supplementary information, acknowledgements, peer review information; details of author contributions and competing interests; and statements of data and code availability are available at https://doi.org/xxxxxxxxxx

## Methods

### Tissue collection and high molecular weight DNA extraction

For extraction of high molecular weight DNA, young leaves were collected from 21-day-old light-grown seedlings. Prior to tissue collection, seedlings were etiolated in complete darkness for 48 h. Flash-frozen plant tissue was ground using a mortar and pestle and extracted in four volumes of ice-cold extraction buffer 1 (0.4 M sucrose, 10 mM Tris-HCl pH 8, 10 mM MgCl2, and 5 mM 2-mercaptoethanol). Extracts were briefly vortexed, incubated on ice for 15 min, and filtered twice through a single layer of Miracloth (Millipore Sigma). Filtrates were centrifuged at 4000 rpm for 20 min at 4°C, and pellets were gently resuspended in 1 ml of extraction buffer 2 (0.25 M sucrose, 10 mM Tris-HCl pH 8, 10 mM MgCl2, 1% Triton X-100, and 5 mM 2-mercaptoethanol). Crude nuclear pellets were collected by centrifugation at 12,000g for 10 min at 4°C and washed by resuspension in 1 ml of extraction buffer 2 followed by centrifugation at 12,000g for 10 min at 4°C. Nuclear pellets were resuspended in 500 ml of extraction buffer 3 (1.7 M sucrose, 10 mM Tris-HCl pH 8, 0.15% Triton X-100, 2 mM MgCl2, and 5 mM 2-mercaptoethanol), layered over 500 ml extraction buffer 3, and centrifuged for 30 min at 16,000g at 4°C. The nuclei were resuspended in 2.5 ml of nuclei lysis buffer (0.2 M Tris pH 7.5, 2 M NaCl, 50 mM EDTA, and 55 mM CTAB) and 1 ml of 5% Sarkosyl solution and incubated at 60°C for 30 min.

To extract DNA, nuclear extracts were gently mixed with 8.5 ml of chloroform/isoamyl alcohol solution (24:1) and slowly rotated for 15 min. After centrifugation at 4000 rpm for 20 min, 3 ml of aqueous phase was transferred to new tubes and mixed with 300 ml of 3 M NaOAc and 6.6 ml of ice-cold ethanol. Precipitated DNA strands were transferred to new 1.5 ml tubes and washed twice with ice-cold 80% ethanol. Dried DNA strands were dissolved in 100 ml of elution buffer (10 mM Tris-HCl, pH 8.5) overnight at 4°C. Quality, quantity, and molecular size of DNA samples were assessed using Nanodrop (Thermofisher), Qubit (Thermofisher), and pulsed-field gel electrophoresis (CHEF Mapper XA System, Biorad) according to the manufacturer’s instructions.

### Tissue collection, RNA extraction and quantification

All tissues were collected in 3-4 biological replicates from different greenhouse-grown plants at approximately 09:00-10:00 AM and flash frozen in liquid nitrogen in 1.5 mL microfuge tubes containing a 5/32 inch (∼3.97 mm) 440 stainless steel ball bearing (BC Precision, TN, USA). Tubes containing tissue were placed in a –80°C stainless steel tube rack and ground using a SPEX^TM^ SamplePrep 2010 Geno/Grinder^TM^ (Cole-Parmer, NJ, USA) for 1 min at 1440 rpm. For shoot apices, total RNA was extracted using TRIzol (Invitrogen, MA, USA) according to the manufacturer’s instructions for ground tissue. For all other tissues (cotyledons, hypocotyls, leaves, flower buds, and flowers), total RNA was extracted using Quick-RNA MicroPrep Kit (Zymo Research). RNA was treated with DNase I (Zymo Research, CA, USA) according to the manufacturer’s instructions. Purity and concentration of the resulting total RNA was assessed using a NanoDrop One spectrophotometer (Fisher Scientific, MA, USA). Libraries for RNA-sequencing were prepared by KAPA mRNA HyperPrep Kit (Roche, Basel, Switzerland). Paired-end 100-base sequencing was conducted on the NextSeq 2000 P3 sequencing platform (Illumina, CA, USA). Reads were trimmed using trimmomatic v0.39^73^ and then mapped to their respective genome using STAR v2.7.5c^74^ and expression computed in transcripts per million (TPM).

### Genome assembly

Reference quality genome assemblies for each of the 22 species (and two reference quality genomes for *S. muricatum*) (**Supplementary Table 2** for accession information) were generated using a combination of long-read sequencing (Pacific Biosciences, CA, USA) for contigging and optical mapping (Bionano Genomics, CA, USA) for scaffolding. Between 1-4 PacBio Sequel IIe flow cells (Pacific Biosciences, CA, USA) were used for the sequencing of each sample (average read N50 = 11,221 bp, average coverage = 53X, average read QV = 83.28). Prior to assembly, we counted k-mers from raw reads with KMC3^75^ (version 3.2.1) and estimated genome size, sequencing coverage, and heterozygosity with GenomeScope2.0^76^. For 5 samples (**Supplementary Table 2** for details), low quality reads were filtered out with a custom script (<u>github.com/pan-sol/pan-sol-pipelines</u>). Sequencing reads from each sample were assembled with hifiasm^77^ exact parameters and software version varied between samples based on the level of estimated heterozygosity and are reported in **Supplementary Table 2**. Post assembly, the draft contigs were screened for possible microbial contamination as previously described^19^.

### Genome assembly scaffolding

Optical mapping (Bionano Genomics, CA, USA) was performed for 17 samples to facilitate scaffolding. Scaffolding with optical maps was performed using the Bionano solve Hybrid Scaffold pipeline with the recommended default parameters (https://bionano.com/software-downloads/). Hybrid scaffold N50s ranged from 33,254,022 bp to 219,385,699 bp (see **Supplementary Table 2** for more detail including Bionano molecules per sample). High-throughput chromosome conformation capture (Hi-C) from Arima Genomics, CA, USA was performed for 8 samples to finalize scaffolding. With Hi-C, reads were integrated with the Juicer (v0.7.17-r1198-dirty) pipeline. Next, misjoins and chromosomal boundaries were manually curated in the Juicebox (v1.11.08) application. Chromosomes were named based on sequence homology, determined with RagTag^78^ scaffold (v2.1.0, default parameters), with the phylogenetically-closest finished genome (see **Supplementary Table 2** for details), 12 of these samples (including nine *S. aethiopicum* samples) were scaffolded with Ragtag. Finally, small contigs (< 50,000 bp) with > 95% of the sequence mapping to a named chromosome were removed. Additionally, small contigs (< 100,000 bp) with > 80% of the sequence mapping to a named chromosome that contained one or more duplicated BUSCO genes, but no single BUSCO genes, were also removed using a python script. Using merqury^79^ with the HiFi data, the final consensus quality of the assemblies was estimated as QV=51.1333 on average and a completeness of 99.2741% on average.

### Gene Annotation

The gene annotation pipeline (**Extended Data Fig. 1d**) involved several crucial steps. Initially, the quality of raw RNASeq reads underwent assessment using FastQC v0.11.9 (http://www.bioinformatics.babraham.ac.uk/projects/fastqc/). Subsequently, reference-based transcripts were generated employing STAR v2.7.5c^74^ and Stringtie2 v2.1.2^80^ workflows. To refine the data, invalid splice junctions from the STAR aligner were filtered out utilizing Portcullis v1.2.0^81^. Orthologs with coverage above 50% and 75% identity were lifted from Heinz v4.0^82^ and Eggplant v4.1^83^ via Liftoff v1.6.3^84^ using parameters –-copies,--exclude_partial and employing both Gmap version 2020-10-14^85^ and Minimap2 v2.17-r941^86^ aligners. In addition, protein evidence from several published Solanaceae genomes^82,83,87^, and the UniProt/SwissProt database were utilized to support gene annotation. Structural gene annotations were generated through the Mikado v2.0rc2^88^ framework, leveraging evidence from the Daijin pipeline. Additionally, microsynteny and orthology to Heinz v4.0 and Eggplant v4.0 were assessed using Microsynteny and Orthofinder v2.5.2^89^. Correction of gene models with inframe stop codons utilized Miniprot2^90^ protein alignments from Heinz v4.0 and Eggplant v4.1. Furthermore, gene models lacking start or stop codons were adjusted by placing them within 300 base pairs of the nearest codon location using a custom python script (<u>github.com/pan-sol/pan-sol-pipelines</u>) (**Supplementary Table 3**). Overall gene synteny was visualized using GENESPACE (v1.3.1)^91^.

For functional annotation, ENTAP v0.10.8^92^ integrated data from diverse databases such as PLAZA dicots (5.0)^93^, Uniprot/Swissprot^94^, TREMBL, RefSeq, Solanaceae proteins, and InterProScan5^95^ with Pfam, TIGRFAM, Gene Ontology, and TRAPID^96^ annotations. Finally, the annotated data underwent a series of filtering steps, excluding proteins shorter than 20 amino acids, those exceeding 20 times the length of functional orthologs and transposable element genes, which were removed using the TEsorter^97^ pipeline.

We assessed the completeness of the gene models by assessing single-copy orthologs through BUSCO^98^ in protein mode, comparing them against the solanales_odb10 *database.* Additionally, we examined the presence or absence of a curated set of 180 candidate genes known to be crucial in QTL studies.

### Transposable element annotation

The *S. lycopersicum* chloroplast and mitochondrion sequences were collected from NCBI reference sequences NC_007898.3 and NC_035963.1, respectively. Non-transposable element repeat sequences including 18S rDNA (OK073663.1), 5S rDNA (X55697.1), 5.8S rDNA (X52265.1), 25S rDNA (OK073662.1), DNA spacer (AY366528.1), centromeric repeat (JA176199.1), and telomere sequences (TTTAGGG) were collected from NCBI and further curated. Transposable element sequences curated in the SUN locus study^99^ as well as several other transposable element sequences from NCBI were also collected. These sequences were combined as the curated set of tomato repeats.

*De novo* transposable element annotation was first performed on each genome using EDTA v2.1.5^100^, with coding sequences from the ITAG4.0 Eggplant V4 annotation^101^ provided (--cds) to purge gene coding sequences in the transposable element annotation and parameters of –-anno 1 –-sensitive 1 for sensitive detection and annotation of repeat sequences. Curated tomato repeats were supplied to EDTA (--curatedlib) for the *de novo* annotation. Transposable element annotations of individual genomes were together processed by panEDTA^102^ for the creation of consistent pan-genome transposable element annotation. Summary of whole-genome repeat annotations were derived from .sum files generated by panEDTA (**Supplementary Table 4**).

Evaluation of repeat assembly quality was performed using LAI b3.2^103^ with inputs generated by EDTA and parameters –t 48 –unlock. LAI of *S. aethiopicum* genomes were standardized based on the HiFi-based reference assembly, with parameters –iden 95.71 –totLTR 49.22 –genome_size 1102623763 –t 48 –unlock.

### Generation of CRISPR-Cas9-induced mutants

CRISPR guide RNAs to target *CLV3* and *SCPL25* across *Solanum* species were designed using Geneious. The Golden Gate cloning approach as described in (29) was used to create multiplexed gRNA constructs. Plant regeneration and *Agrobacterium tumefaciens*-mediated transformation of *S*. *prinophyllum* were performed according to our previously published protocol^104^. For *S. cleistogamum* plant regeneration, the medium was supplemented with 0.5 mg/L zeatin instead of 2 mg/L and for the selection medium, 75 mg/L kanamycin was used instead of 200 mg/L. For *S. aethiopicum*, the protocol was the same as for *S. cleistogamum*, except the fourth transfer of transformed plantlets is done onto medium supplemented with 50 mg/L kanamycin. Seed germination time in culture can vary between species and batches of harvested seeds. Typically, *S*. *prinophyllum* germination took 8-10 days, *S*. *cleistogamum* germinated in 6-8 days, and *S. aethiopicum* in 7-10 days.

### Distribution maps and species status

Species were categorized into wild, domesticated, locally-important consumed, or ornamental based on taxonomic literature and expert opinion^8^ (PBI *Solanum* Project (2024). *Solanaceae* Source. http://www.solanaceaesource.org/). Native ranges were derived from the same taxonomic literature and approximate centroids of the ranges were used for the mapping.

### Phylogenomic analyses

*Jaltomata sinuosa* was used an outgroup for the *Solanum* pan-genome tree, whereas the closely related *S. anguivi*, *S. insanum*, and *S. melongena* were used as an outgroup for the *Solanum aethiopicum* dataset. Orthofinder^89^ was used to identify single copy orthologs across all species. This resulted in 7,825 loci for the *Solanum* pan-genome dataset, and 19,769 loci for the *S. aethiopicum* dataset. To reduce computing time, we randomly subsampled 5,000 loci for the *S. aethiopicum* dataset. To reduce the effect of missing data and long branch attraction, sequences shorter than 25% of the average length for each loci were eliminated, following Gagnon *et al.* (2022)^23^. MAFFT^105^ was used to align each locus individually. Only loci that had all species in the alignment were kept. trimAl was also used to remove columns that had more than 75% gaps. IQ-TREE2^106^ was used to generate individual ML trees for each locus. The resulting phylogenies were used for coalescent analyses with ASTRAL-III version 5.7.3^107^, where tree nodes with <30% BS values were collapsed using Newick Utilities version 1.5.0^108^. Branch support was assessed using localPP support^109^, where PP values >0.95 were considered strong, 0.75 to 0.94 weak to moderate, and ≤0.74 as unsupported. Trees were visualized with R using the packages ggtree^110^ and treeio^111^.

### Gene expansion contraction analysis

To analyze gene expansions and contractions, we processed the ultrametric species tree and gene family counts from OrthoFinder using CAFE5^112^. CAFE5 was run with the gamma model and parameter ’k=3’ to identify changes in gene family size along the species tree while accounting for rate variation among gene families.

### GO enrichment analysis

Gene Ontology (GO) enrichment analysis was performed using the GOATOOLS package^113^ to investigate the functional implications of genes associated with various duplication types including whole-genome (WGD), tandem (TD), proximal (PD), transposed (TSD) and dispersed (DSD) duplications. Genes were classified into these different duplication categories by DupGenefinder^30^. Additionally, we conducted GO enrichment on gene expansions (**Supplementary Table 5**) and contractions (**Supplementary Table 6**) identified across all lineages as reported by CAFE5, to explore functional trends related to these gene copy number changes across the pangenome.

### Synteny analysis

Genomic neighborhood around *CLV3* for selected species was manually inspected to detect and annotate intact and pseudogenized *CLV3* copies using pairwise sequence comparison with Exonerate (www.ebi.ac.uk/about/vertebrate-genomics/software/exonerate). Synteny plots were generated from a reciprocal BLASTP table obtained running Clinker (v0.0.29, <u>github.com/gamcil/clinker</u>). Pseudomolecule visualization was generated *via* a custom script (<u>github.com/pan-sol/pan-sol-pipelines</u>). Transposable elements and resistance genes annotations were overlaid as needed using custom scripts (<u>github.com/pan-sol/pan-sol-pipelines</u>).

### Gene expression analysis

Reads from each tissue sample were aligned to the corresponding species-specific genome using STAR v2.7.2b^74^, and only samples with more than 50% uniquely mapped reads were retained for subsequent analysis. For each species with two or more biological replicates per tissue, we calculated the Spearman correlation between tissue replicates, and removed samples with low correlation (0.75 or below). This yielded gene expression estimates for 271 samples across 22 species, with 15 species having expression data in two or more tissues. Expression data was TPM-normalized and genes with zero expression across all samples were excluded from further analysis. Principal component analysis was performed on the tissue-specific expression profiles of 5,146 singleton genes shared across all 22 species to reveal the global relationships among samples.

#### Is the total dosage of duplicate gene pairs conserved across Solanum?

Survival of a gene after duplication depends on the competition between preservation to maintain partial or total dosage and mutational degradation rendering one copy with reduced or no function. Consequently, functional fates of duplicate genes are often characterized by the extent of selective pressures on total dosage. To assess the relative importance of dosage balance (copies evolving under strong purifying selection to maintain total dosage) and neutral drift (no selection on total dosage) in maintaining duplicate genes, we compared the total expression of paralog pairs within each tissue for each pair of species. Note that the prickle tissue from *S. prinophyllum* is not included in this analysis since it is absent in the other 21 species.

In each tissue, gene expression was averaged over the biological replicates for each species. For each pair of species with expression data in a shared tissue, orthogroups with exactly two copies in each species with non-zero average expression in the tissue were retained for further analysis. For each tissue and species pair, we calculated the summed expression of paralog pairs in each retained orthogroup, and observed that the total “orthogroup-level” expression was highly correlated across species suggesting a prominent role of dosage balance in shaping the expression evolution of paralogs. We computed the ratio of the orthogroup-level expression between the species pair and transformed them into z-scores. For each orthogroup in a species expressed in the tissue of interest, we averaged the *p*-values from all pairwise species comparisons, adjusted the average *p*-values using Benjamini-Hochberg correction, and classified orthogroups with adjusted average *p*-value < 0.05 as dosage-unconstrained orthogroups. All other orthogroups in the species and tissue were assumed to be evolving under constraint on total dosage.

All other orthogroups were assumed to evolve under selective constraint on total dosage. Note that the high z-score threshold provides a conservative estimate of the number of paralog pairs evolving under drift. Sequence evolution rates for paralog pairs (Ka/Ks) were calculated using KaKs_Calculator 2.0^114^.

#### Different modes of paralog functional evolution

For each of the 15 species with expression in two or more tissues, the expression data was first subset to genes with more-than-median expression in at least one sample. Coexpression network for each species was constructed by calculating the Pearson correlation between all pairs of genes, ranking the correlation coefficients for each gene (with NAs assigned the median rank), and then standardizing the network by the maximum ranked correlation coefficient. Coexpression for each pair of paralogs in each orthogroup was obtained from this rank-standardized network. For each paralog pair with non-zero expression in two or more samples, we also computed the fold-change of expression across samples and used the absolute values of mean and standard deviation (SD) of log2-transformed fold-change across samples to summarize the degree of expression divergence between the two copies.

We classified the paralog pairs within each species into different retention categories based on their variation in expression levels and correlated expression across samples. We selected these two axes of variation since they intuitively represent average expression difference (fold-change) and specific pattern of difference (coexpression) between gene pairs. We classified paralog pairs into four broad groups as follows:

I. Dosage-balanced: coexpression > 0.9, mean log_2_ fold-change < 1, SD of log_2_ fold-change < 1
II. Paralog dominance: coexpression > 0.9, mean log_2_ fold-change >= 1, SD of log_2_ fold-change < 1
III. Specialized: coexpression > 0.9, mean log_2_ fold-change >= 1, SD of log_2_ fold-change >= 1
IV. Diverged: coexpression < 0.5, mean log_2_ fold-change >= 1, SD of log_2_ fold-change >= 1

Paralogs originating from whole genome (WGD), tandem and proximal duplications were obtained using the DupGen_finder pipeline^30^. WGD pairs with *Ks* ranging from 0.2 to 2.5, and tandem and proximal duplicates with *Ks* ranging from 0.05 to 2.5 were used to generate the stacked bar plots corresponding to whole genome and small-scale duplications, respectively, in **Fig. 3i**.

Gene family size for each classified paralog pair within a species corresponds to the number of genes in its orthogroup. The expression breadth of a gene corresponds to the number of tissues (among apices, cotyledon, hypocotyl, inflorescence, leaves) where the gene has an average expression greater than 3 TPM. Number of shared tissues expressing a paralog pair is computed by intersecting the expression breadths of both copies, and ranges from 0 to 5. A gene was considered non-functional if it was annotated as a pseudogene or had an average expression below 3 TPM. Tissue-specific genes for each tissue were identified as genes with the highest expression in the tissue of interest, tissue-specificity score^115^ greater than 0.7 and with expression greater than 5 TPM in the relevant tissue.

### Mapping of loci controlling *S. aethiopicum* locule number

The high-locule count parent and reference accession PI 424860, and low-and higher-locule count parents 804750187 and 804750136, respectively, were selected as founding parents to map QTLs and their causative variants affecting fruit locule number. Resulting F1 progeny were selfed to generate F2 mapping populations, which were sown in the greenhouse and then transplanted to a field site at Lloyd Harbor, New York, USA, during the summer of 2022. Approximately 10 fruits were collected from each F2 individual and the number of locules exposed by slicing the fruit transversely and counting. In the 804750187 x PI 424860 and 804750136 x PI 424860 derived F2 populations, 144 and 135 individuals were phenotyped, respectively. For each population, DNA from 30 random individuals at the low and high ends of the phenotypic distribution for locule number were pooled for bulk-segregant QTL-Seq analysis. The DNA from 8 individuals of the common parental accession PI 424860 were also pooled to capture parental polymorphisms.

DNA from 30 randomly selected low-and high-locule count individuals was extracted from young leaf tissue using a DNeasy Plant Pro Kit (Qiagen, Hilden, Germany) according to the manufacturer’s instructions for high-polysaccharide content plant tissue. Tissue used for extraction was ground using a SPEX^TM^ SamplePrep 2010 Geno/Grinder^TM^ (Cole-Parmer, NJ, USA) for 2 min at 1440 rpm. Sample DNA (1 µL assay volume) concentrations were assayed using Qubit 1X dsDNA HS buffer (ThermoFisher, MA, USA) on a Qubit 4 fluorometer (ThermoFisher, MA, USA) according to the manufacturer’s instructions. Separate pools were made for the parents, the bulked high-locule count F2 individuals, and the bulked low-locule count F2 individuals, with an equivalent mass of DNA pooled from each individual to yield a final pooled mass of 3 µg in each bulk. DNA pools were purified using 1.8X volume of AMPure XP beads (Beckman Coulter, CA, USA) and the DNA concentration and purity assayed by Qubit and a NanoDrop One spectrophotometer (Fisher Scientific, MA, USA), respectively.

Paired-end sequencing libraries for QTL-Seq analysis were prepared with >1 µg of DNA using a KAPA HyperPrep PCR-free kit (Roche, Basel, Switzerland) according to the manufacturer’s instructions. Indexed libraries were pooled for sequencing on a NextSeq 2000 P3 chip (Illumina, CA, USA). Mapping was performed using the end-to-end pipeline implemented in the QTL-Seq software package^116^ (v2.2.4, <u>github.com/YuSugihara/QTL-seq</u>) with reads aligned against the *S. aethiopicum* (Saet3, PI 424860) genome assembly.

To determine the effects of the two identified QTL on locule number in the 804750136 x PI 424860 derived populations, co-segregation analysis was performed on the full F2 populations by genotyping *SaetCLV3* and the minor effect locus on chromosome 5. For *SaetCLV3*, a cleaved amplified polymorphic sequence (CAPS) assay was used to genotype a variant in the promoter region of *SaetCLV3* linked to the identified *CLV3* SV haplotypes. A 1258 bp region surrounding an *Ase*I restriction fragment length polymorphism (RFLP) in the *SaetCLV3* promoter was amplified using KOD One^TM^ PCR Master Mix (Toyobo, Osaka, Japan) on template DNA extracted by the cetyltrimethylammonium bromide (CTAB) method^117^ (see **Supplementary Table 13** for primers 5431 & 4681). To 5 µL of the resulting PCR product, a 10 µL reaction containing 0.2 µL *Ase*I (New England BioLabs, MA, USA) and 1 µL CutSmart^TM^ r3.1 Buffer (New England BioLabs, MA, USA) was incubated for 2 hours at 37 °C. The reactions were then loaded onto a 1% agarose gel and electrophoresed in an Owl^TM^ D3-14 electrophoresis box (Thermo Scientific, MA, USA) containing 1X TBE buffer for 30 min at 180 V delivered from an Owl^TM^ EC 300 XL power supply (Thermo Scientific, MA, USA). The electrophoresis results were visualized under UV light using a Bio-Rad ChemiDoc^TM^ XRS+ (Bio-Rad, CA, USA) imaging platform and ImageLab^TM^ (Bio-Rad, CA, USA) software. Resulting banding patterns were then used to assign genotypes. For the chromosome 5 QTL, primers (see **Supplementary Table 13** for primers 5883 & 5884) were used to amplify a 425 bp region harboring a 1 bp deletion occurring near the summit of the QTL peak using KOD One^TM^ PCR Master Mix. The resulting PCR products were purified using Ampure 1.8X beads and served as template for Sanger sequencing (Azenta Genewiz, NJ, USA). The sequencing results were then used to assign genotype calls at chromosome 5.

### Conservatory analysis

The Conservatory algorithm (V2.0)^36^ was employed to identify conserved noncoding sequences (CNSs) within the *Solanaceae* family (**Extended Data Figure 2d**) (https://conservatorycns.com/dist/pages/conservatory/about.php). A total of 26 genomes, including 23 *Solanum* genomes, two tomato genomes (Heinz and M82) and one groundcherry (*Physalis grisea*), were used as references to enable the identification of CNSs irrespective of structural variations among references. Protein similarity was scored using Bitscore^118^, while *cis*-regulatory similarity was assessed using LastZ^119^ score. Homologous gene pairs were required to share at least one CNS. For orthogroup calling, all orthologous genes shared at least one CNS with the reference gene. Gene pairs with a conservation score exceeding 90% of the highest score were classified as paralogs (**Extended Data Figure 2b**). A total of 844,525 paralogs were identified across the *Solanum* pan-genome. Sequence evolution pressure rates (Ka/Ks) for paralog pairs were calculated using the R seqinR package (v4.2-36)^120^. Gene duplication events were classified using DupGenefinder^30^, identifying whole-genome (WGD) and transposed (TSD) duplications for gene pairs recognized by both Conservatory and DupGenefinder tools. Tandem (TD) and proximal (PD) duplications were defined based on gene positioning: adjacent genes were considered TD, and genes up to 10 genes apart were defined as PD. All other duplicated gene pairs were categorized as dispersed (DSD) duplications (**Extended Data Figure 2c**). Of the identified paralogs, 23,730 were associated with expression groups and were used to compare relationships between sequence evolution pressure rates and protein and *cis*-regulatory divergence across different expression groups. Homologs, orthogroups, and paragroups were identified, and relationships between protein and *cis*-regulatory elements were visualized using custom scripts, which are available on GitHub (<u>github.com/pan-sol/pan-sol-pipelines</u>).

### Statistical analysis

All statistical tests were performed in R. For the quantitative analysis of fruit locule numbers in Figures 4f, 6c, 6d, and Extended Data Figure 6b, n represents the “number of fruits quantified.” Pairwise comparisons were conducted using Dunnett’s T3 test for multiple comparisons with unequal variances, with default parameters (see **Supplementary Tables 14-17**).

### Reporting summary

Further information on research design is available in the Nature Portfolio Reporting Summary linked to this article.

### Data availability

All data are available within this Article and its Supplementary Information. Raw sequencing data are available in the SRA under BioProject PRJNA1073673. Genome, expression, and phenotypic data are available at Solpangenomics website (www.solpangenomics.com). Paralog expression analysis scripts are available at <u>github.com/gillislab/pansol_expression_analysis</u>. Other analysis scripts are available within <u>github.com/pan-sol/pan-sol-pipelines</u>.

## Acknowledgements

We thank members of the Lippman laboratory and critical friends M. Bartlett, Y. Eshed, and I. Efroni for discussions and feedback. We thanks B. Seman from the Lippman lab for technical support, and T. Mulligan, K. Schlecht, and S. Qiao for assistance with plant care. We thank E. Cruickshank and T. Jenike for Pepino dulce fruit images. We thank S. Muller, R. Wappel, S. Mavruk-Eskipehlivan, and E. Ghiban from the CSHL Genome Center for sequencing support. MB is grateful for financial support from the Plant Health and Environment department of the French National Institute for Agriculture, Food, and Environment (INRAE). JWS is supported by an NSF Postdoctoral Fellowship in Biology (IOS-2305651). MJP is funded by the William Randolph Hearst Scholarship from the Cold Spring Harbor School of Biological Sciences. SO is supported by the OSU STEM Education Faculty Startup Award and the Global Gateways Initiative Grant. Funded by the Convenio 566 of 2014 between Universidad Nacional de Colombia and Minciencias (GPS and FR). UCU Research Funds (EBK). WRM is the Davis Family Professor of Human Genetics. Work on *Solanum* taxonomy, morphology and phylogenetics was funded by NSF Planetary Biodiversity Initiative grant “PBI Solanum: a worldwide initiative” (DEB-0316614 to SK), Sibbald Trust (RH), Fonds de recherche du Québec – Nature et Technologies postdoctoral fellowship (EG), and National Geographic Society Northern Europe Award GEFNE49-12 (TS). National Institutes of Health grant R01MH113005 (JG). National Science Foundation Plant Genome Research Program grant IOS-2216612 (AF, JG, JVE, MCS, ZBL). The Howard Hughes Medical Institute (ZBL). Finally, we thank the indigenous peoples of Australia on whose ancestral lands *S. cleistogamum* and *S. prinophyllum* grow.

## Author Contributions

Conceptualization: MCS, ZBL

Data curation: MB, KMJ, JWS, SR, AH, MJP, HS1, MA, XW, SO

Formal analysis: MB, KMJ, JWS, SR, IG, AH, MJP, HS1, MA, XW, SO, EG, TS, AF, JG, JVE, MCS, ZBL

Funding acquisition: MB, JWS, MJP, SO, GPS, FR, EBK, EG, SK, TS, AF, JG, JVE, MCS, ZBL

Investigation: MB, KMJ, JWS, SR, IG, AH, MJP, HS1, HS2, MA, XW, SO, JG, JVE, MCS, ZBL

Methodology: MB, KMJ, JWS, SR, AH, MJP, HS1, GMR, MA, XW, SO, YG, KS, EG, SK, TS, JG, JVE, MCS, ZBL

Project administration: MB, KMJ, AF, JG, JVE, MCS, ZBL

Software: KMJ, SR, AH, MA, SO, MCS

Resources: MB, KMJ, JWS, HS2, BF, MA, XW, RS, JH, HG, YG, KS, GPSR, AO, FR, SG, WRM, EG, SK, TS, AF, MCS

Supervision: JG, JVE, MCS, ZBL

Validation: MB, KMJ, JWS, SR, AH, SO, EBK, EG, SK, TS, AF, JG, JVE, MCS, ZBL Visualization: MB, KMJ, JWS, SR, AH, GMR, EBK, JG, JVE, MCS, ZBL

Writing – original draft: MB, KMJ, MCS, ZBL

Writing – review & editing: MB, KMJ, JWS, SR, IG, AH, MJP, HS1, HS2, SO, EBK, EG, SK, TS, AF, JG, JVE, MCS, ZBL

## Additional information

Supplementary information The online version contains supplementary material available at https://doi.org/xxxxxx

Correspondence and requests for materials should be addressed to Jesse Gillis, Joyce Van Eck, Michael C. Schatz, or Zachary B. Lippman.

## EXTENDED FIGURE LEGENDS

**Extended Data Figure 1: *Solanum* pan-genome species (selected images), *de novo* assemblies, and gene annotation pipeline. (a)** Phenotypic diversity of shoots and fruits (where available) from a subset of the species selected for the *Solanum* pan-genome. Scale bars: 5 cm (shoots) and 1 cm (fruits). **(b)** Total sizes of the pan-*Solanum* genome assemblies evaluated by cumulative sequence length. Genomes of tomato (*S. lycopersicum*, Heinz SL4.0 and M82) and Brinjal eggplant (*S. melongena*, V3) are shown as references. **(c)** Hi-C contact map from *S. candidum* shown as a representative example of data used to generate chromosome-scale assemblies. **(d)** Flow chart depicting the gene annotation pipeline used in this study, noting the required input data (RNA-seq data, protein alignments, and genome sequences), tools, and customs scripts. Preprocessing, annotation, homology, functional annotation, and packaging steps are detailed.

**Extended Data Figure 2: Comparative genomic analysis of orthogroup dynamics and Conservatory analysis of paralogous gene pairs across pan-*Solanum* species. (a)** Functional enrichment for orthogroup expansions and contractions in tomato, eggplant, and major *Solanum* clades. The top five enriched GO terms per species/clade are shown. Circle size represents gene ratio. **(b)** Comparison of orthogroups conservation group size and the subsequent paragroups, defined by the number of species having paralogous genes. Note that ∼60% of duplicated gene orthogroups are conserved across all *Solanum* pan-genome species (Core), while less than 1% of the paragroups are Core. **(c)** Duplicated gene pairs classification of the pan-genome species according to duplication type. **(d)** Flow chart of the Conservatory tool used to define conserved non-coding sequences (CNSs) across pan-genome orthogroups and paragroups. **(e)** Divergence of protein and *cis*-regulatory sequences across increasing evolutionary pressure, as measured by Ka/Ks values, for the indicated types of gene duplications. For each duplication type the predicted mean, residuals, and 0.95 confidence interval of the normalized BLASTP and LastZ scores are shown.

**Extended Data Figure 3: Paralog pairs expression analysis. (a)** Schematic of dosage-constrained and dosage-unconstrained orthogroups reflecting different degrees of selection on the total dosage of paralog pairs across species. Orthogroup 1 has paralog pairs with identical total dosage across species, whereas orthogroup 2 has different total dosages in each species. For each tissue, orthogroup and species, the total dosage of two paralogs is compared with that of the two homologs in each of the remaining species, and deviations from the expected ratio of total dosages are classified as “unconstrained”. This is repeated for all species that share the orthogroup and expressed in the tissue of interest, and the majority classification across species is taken as the classification for the entire orthogroup. Therefore, orthogroup 1 is classified as “dosage-constrained” while orthogroup 2 is classified as “dosage-unconstrained”. **(b)** The fraction of uniquely mapped reads for each tissue sample and species (left), and the average gene expression correlation with other samples from the same tissue and species (right). Red arrows in both cases point to the five outlier samples excluded from further analysis. **(c)** Sankey plot shows the concordance between classification of paralog pairs based on two independent approaches (total dosage conservation and conservation of expression levels and profiles). Thickness of lines connecting each pair of groups shows the odds ratio of enrichment. **(d)** Line plots showing examples of paralog pairs in each of the four groups of paralog expression patterns. **(e)** Functional enrichment for paralog pairs from the different groups. The top five enriched GO terms per expression group is shown. Circle size represents gene ratio. **(f)** Relationship of protein and *cis*-regulatory sequence conservation on the different paralog expression groups over increasing evolutionary pressure. For each expression group the predicted mean, 95% confidence interval, and residuals of the normalized LastZ score are shown.

**Extended Data Figure 4: Extreme variation in transposable elements and resistant gene content at the *CLV3* locus across *Solanum*. (a)** Gene and transposable element compositions are highly variable at the *CLV3* locus across the eggplant clade. While most of the gene content shows collinearity, the transposable element profile and density varies considerably. Stacked bars show the absolute number and type of transposable element for the window of three genes. **(b)** Microsyntenic relationships at the *CLV3* locus across the eggplant clade show dynamic expansions and contractions of resistance genes. Resistance genes are identified by blue dots. Presence-absence of *CLV3* paralogs is shown as in Figure 4. Lineage-specific *CLV3* duplications denoted with asterisks. Window sizes range from 397,829 bp (*S. torvum*) to 634,079 bp (*S. aethiopicum*) and are centered on the *CLV3* locus. Functional *CLV3* copies are denoted by full circles while truncated/pseudogenized copies are shown as half circles, as in Figure 4. Grey lines illustrate conservation, while blue lines represent loss of synteny. **(c)** CRISPR/Cas9 gene-edited loss-of-function null alleles of *CLV3* genes in *S. prinophyllum* and *S. cleistogamum*. **(d)** CRISPR/Cas9 gene-edited loss-of-function null alleles of African eggplant *SaetCLV3a/b*. Numbers represent the proportion of cloned and sequenced *SaetCLV3a/b* alleles as a ratio of the total number of clones sequenced in the three first-generation transgenic (T0) plants showing fasciation phenotypes.

**Extended Data Figure 5: Structural variants and gene copy number variation in the African eggplant pan-genome. (a)** Structural variant density across all chromosomes in African eggplant and its wild progenitor *S. anguivi* in 2 Mbp windows. **(b)** Percentage of structural variants overlapping with different genomic features. **(c)** Gene presence-absence and copy number variation in 17 orthogroups containing known genes regulating three major domestication traits in tomato across the African eggplant pan-genome and *S. anguivi*. Stars mark gene truncation or pseudogenization.

**Extended Data Figure 6: Interactions between the *CLV3* and Chr5 African eggplant locule number QTLs in F2 populations**. **(a)** Averaged fruit locule number counts for plants from the 804750136 x PI 424860 (top) and 804750187 x PI 424860 (bottom) segregating F2 populations. Average locule counts for the parental genotypes are also shown. **(b)** Stacked bar plots showing fruit locule number from ranked F2 population-derived genotypes segregating the reference (REF) and alternative (ALT) alleles of *SaetCLV3* and the chromosome 5 QTLs. P: parents.

