## Supplementary figures and images for "*Solanum* pan-genomics and pan-genetics reveal paralogs as contingencies in crop engineering"

### Extended Figure 1

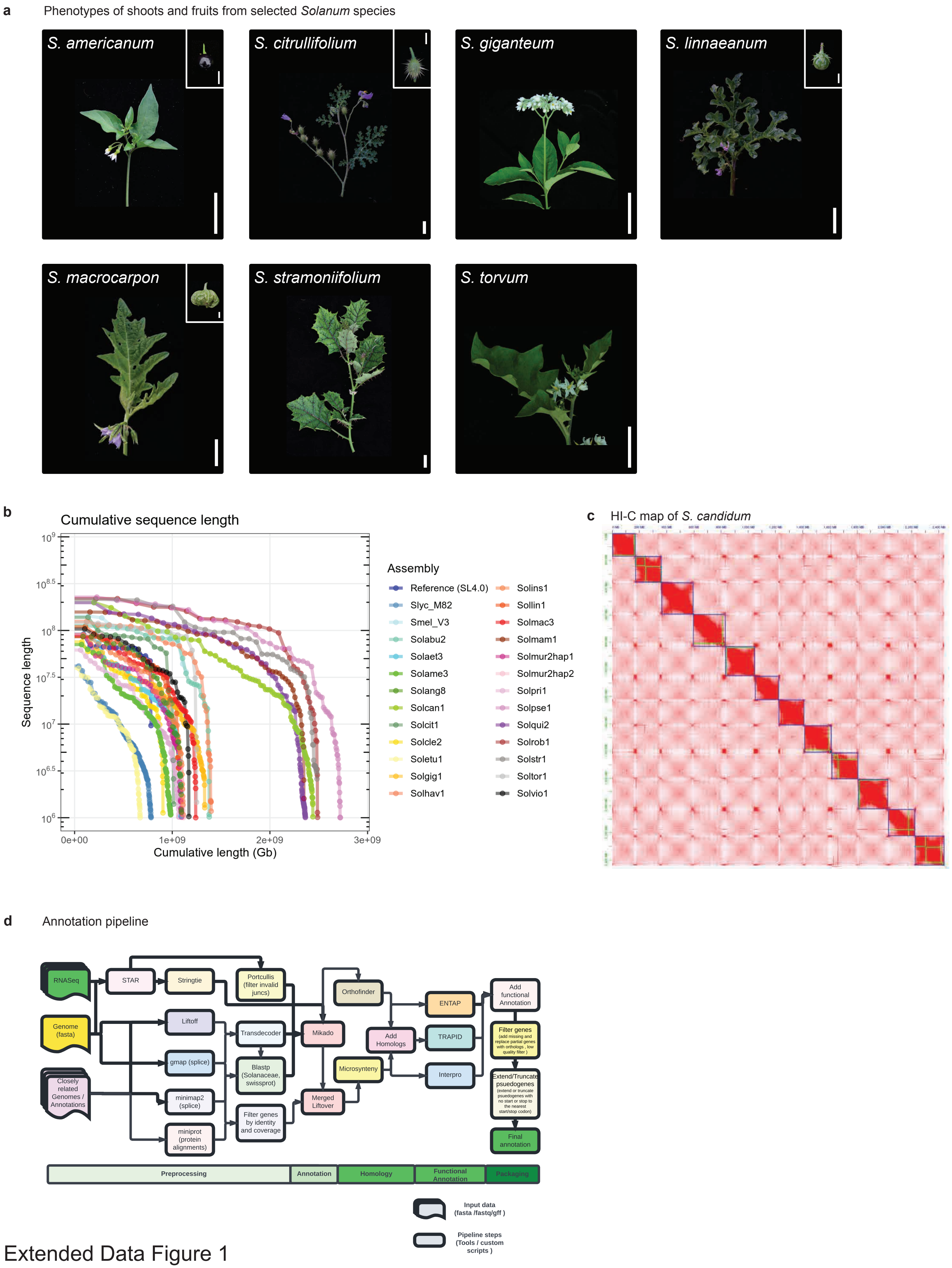


Extended Data Figure 1

### Extended Figure 2

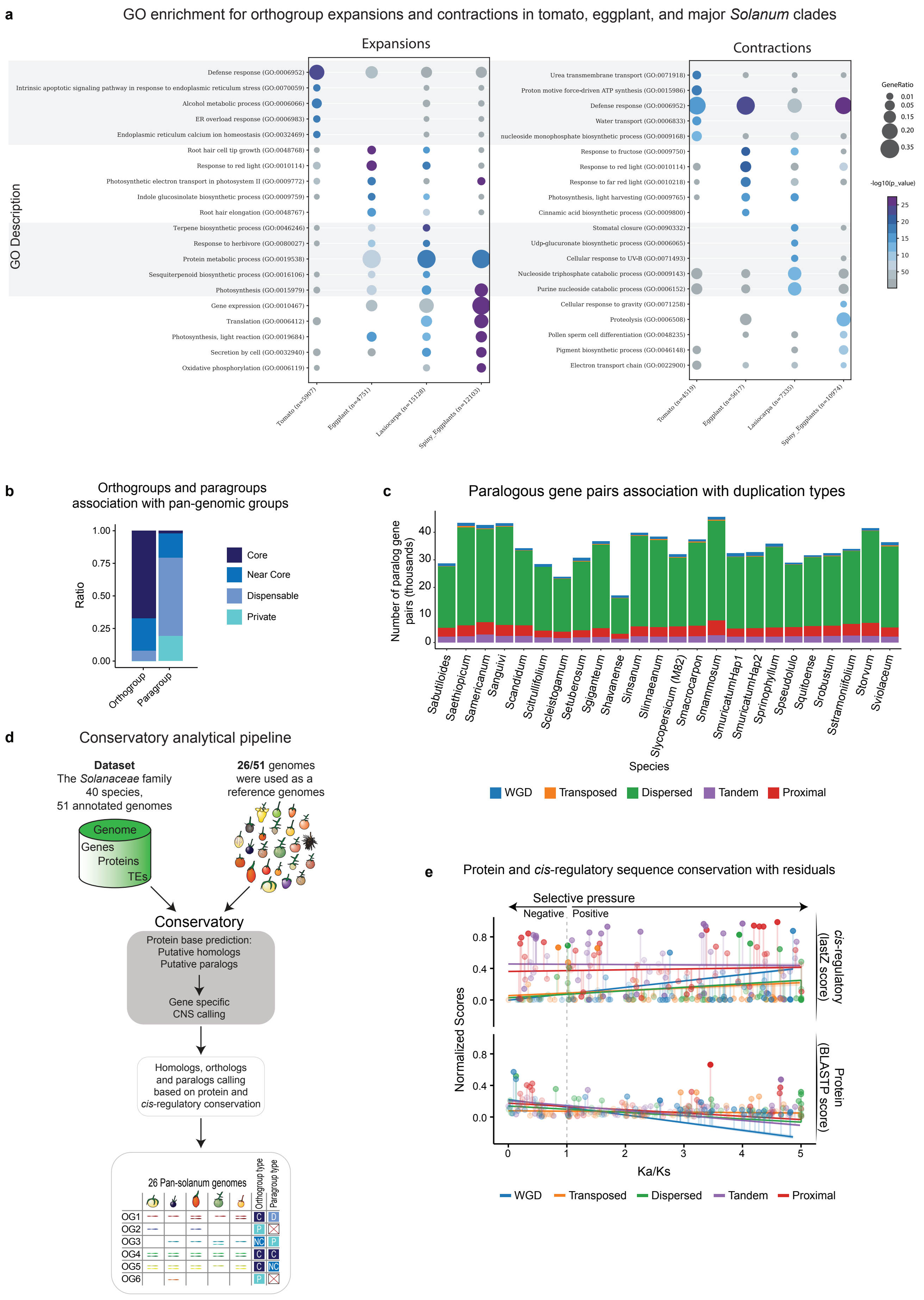


Extended Data Figure 2

### Extended Figure 3

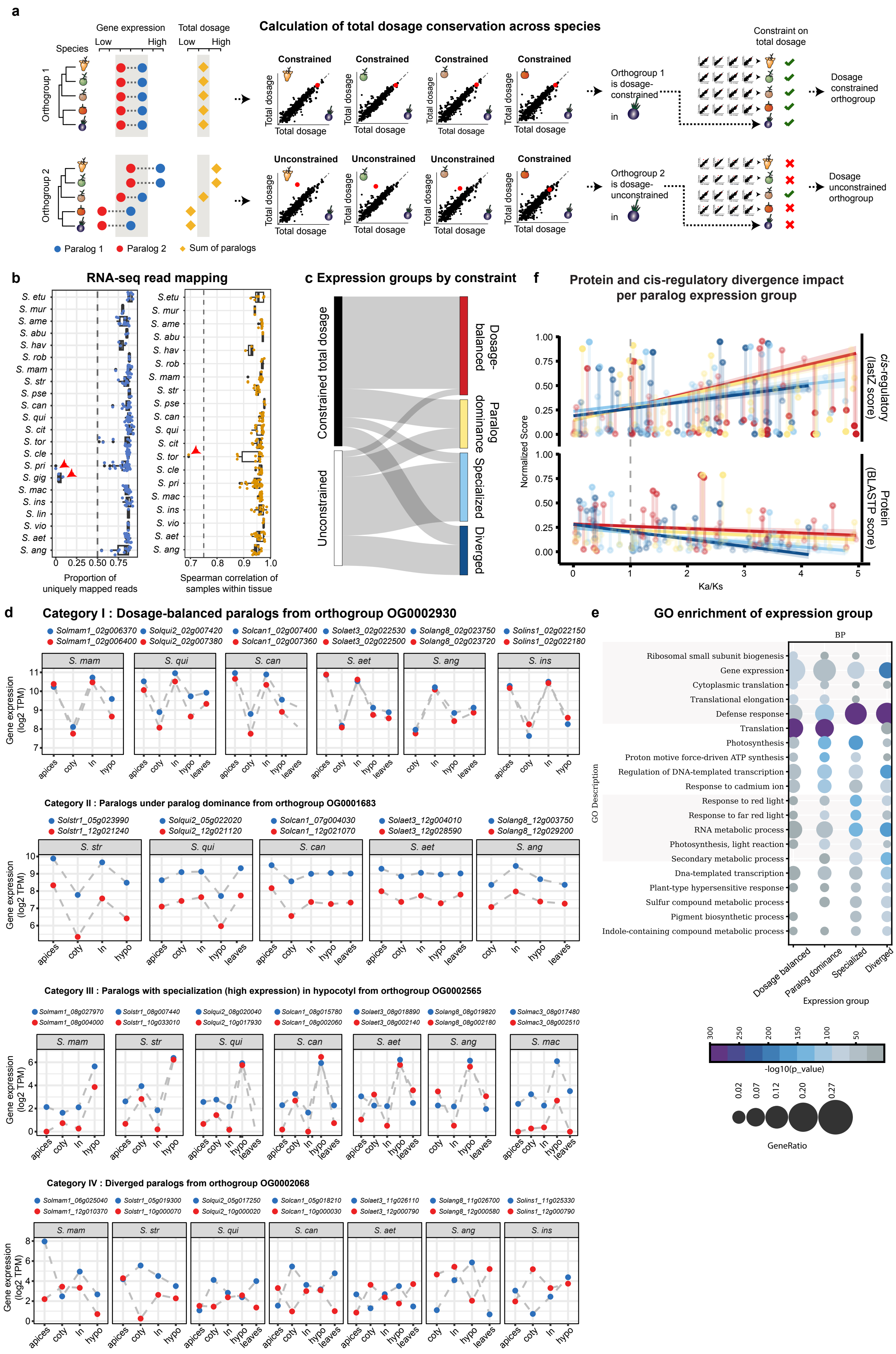


Extended Data Figure 3

### Extended Figure 4

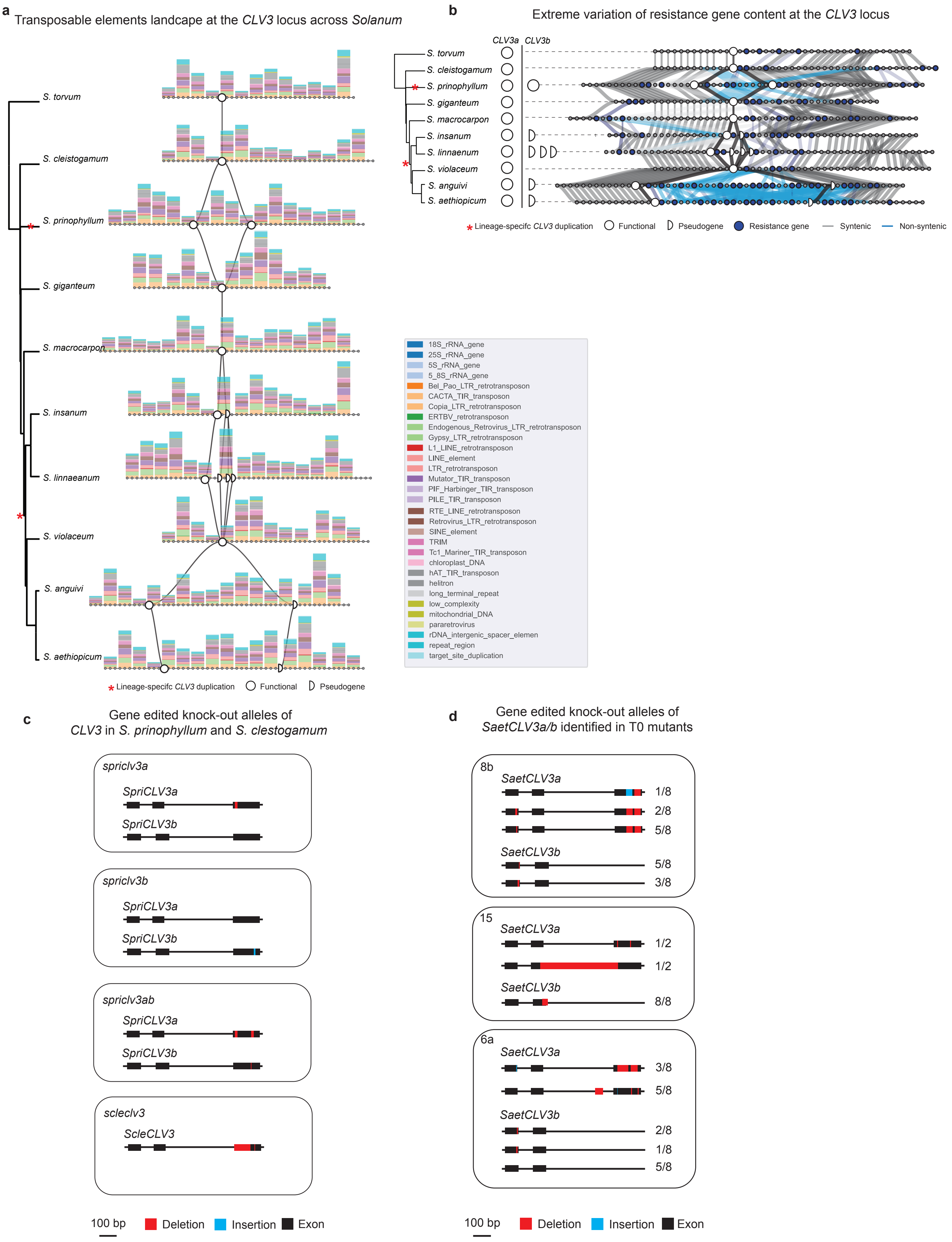


Extended Data Figure 4
