## Extended Figure 5 for "*Solanum* pan-genomics and pan-genetics reveal paralogs as contingencies in crop engineering"

**a** SV density across all chromosomes in African eggplant and *S. anguivi*

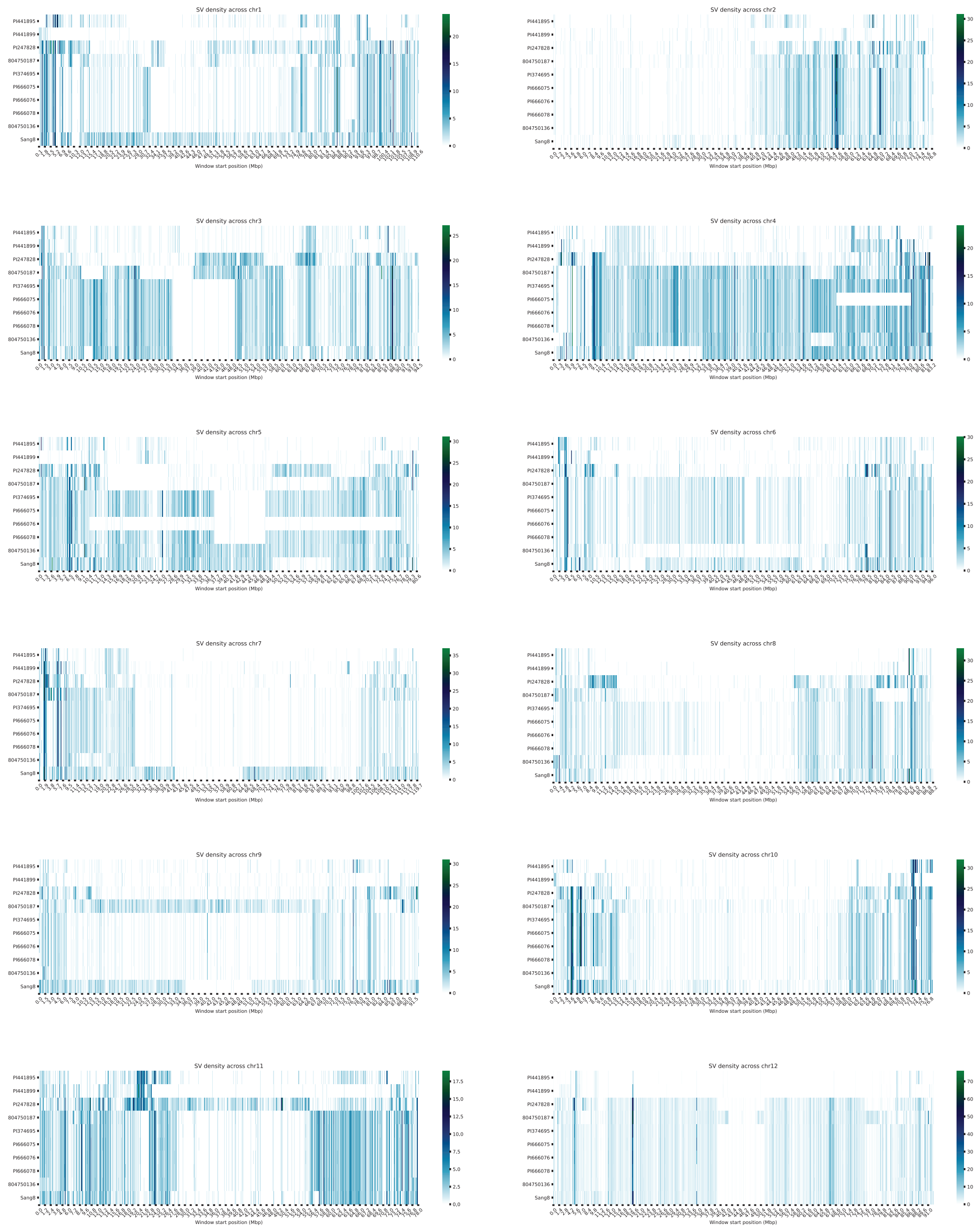

**b** Percentage of SVs overlapping with various genomic features in African eggplant

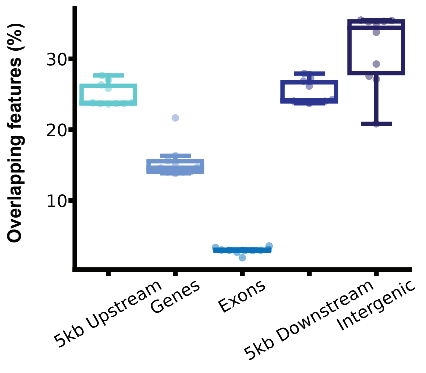


**c** Variation in developmental genes and their paralogs underlying major domestication traits

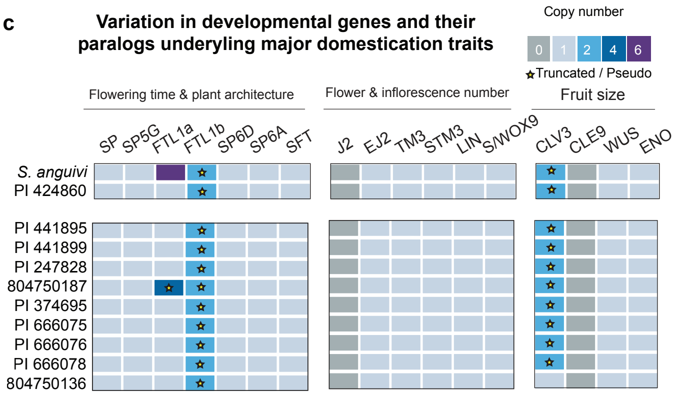


Extended Data Figure 5
