## Extended Figure 6 for "*Solanum* pan-genomics and pan-genetics reveal paralogs as contingencies in crop engineering"

**a** **Locule number count for segregating F2 populations**

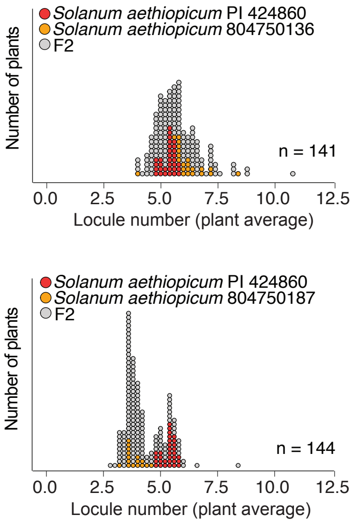


**b** **Locule number count for all allelic combinations**

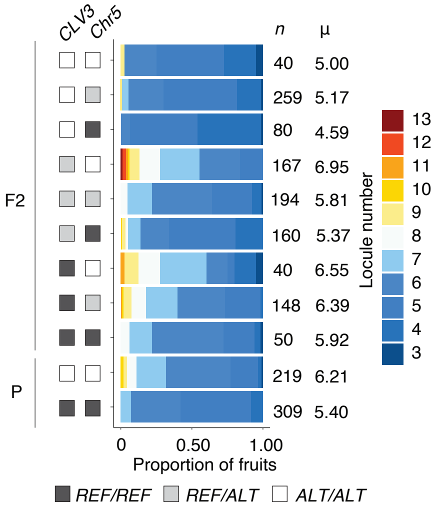


Extended Data Figure 6
